# HDOCK-Multimer: integrating docking and combinatorial assembly for structure prediction of large protein complexes

**DOI:** 10.64898/2026.08.06.736029

**Authors:** Xuan Yao, Yifan Ya, Hao Li, Sheng-You Huang

## Abstract

Deep learning methods, such as AlphaFold and RosettaFold, achieve high accuracy in protein structure prediction. However, predicting the structure of large protein complexes remains challenging due to their large size and intricate multi-chain interactions. Docking-based methods can handle large proteins, but are limited by the huge combinatorial binding space of multi-chains. Assembly-based approaches offer an alternative, but their accuracy critically relies on the precision of predicted subcomponents. Addressing the challenges, we propose HDOCK-Multimer (HDM), a structure prediction framework of large protein complexes by integrating ab initio docking and combinatorial assembly. HDM can efficiently reduce reliance on subcomponent accuracy through docking process, while leveraging the pairwise interactions of subcomponents through assembly strategy. HDM is extensively validated on three benchmarks of 35 large heteromeric complexes, 172 large protein complexes, and 7 CASP15 targets, and compared with state-of-the-art methods including MoLPC, CombFold, AlphaFold-Multimer (AFM), and AlphaFold3 (AF3). It is shown that HDOCK-Multimer substantially outperforms the other methods. In addition, HDM also shows ability to predict the stoichiometry and model the complex without stoichiometry input. It is anticipated that HDM will serve as a powerful tool for studying large protein complexes or molecular machines. The HDM package is freely available at https://github.com/huang-laboratory/HDOCK-Multimer/.

## 1 Introduction

Proteins exert their functions in the form of complexes within cells^1–3^. Large protein complexes play essential roles in various biological processes, such as transport^4^, signal transduction^5^, and gene regulation^6^. Therefore, determining the structure of protein complexes is critical for understanding their functional mechanisms and related drug discovery. Experimental methods^7, 8^, such as X-ray crystallography, nuclear magnetic resonance (NMR), and cryo-electron microscopy (cryo-EM), face limitations in resolving these structures^9^, including high cost, time consumption, challenges in sample preparation, and resolution constraints. Relying solely on experimental approaches cannot meet the growing demand for the need of structural data. As such, computational methods are necessary for the prediction of protein complex structures^10^.

Traditional computational methods for protein complex structure prediction can be divided into two main categories: template-based and docking-based approaches. Template-based modeling^11–13^ can effectively address interface flexibility by leveraging homologous complex structures but is critically limited by template availability and quality. Docking-based approaches^14–17^, which are initially developed for dimers, face a combinatorial explosion in chain arrangement possibilities when extended to multimeric systems, leading to substantially increased computational cost. Some methods incorporate symmetry constraints to reduce the sampling space and have achieved some successes in cyclic (*C_n_*) symmetry and dihedral (*D_n_*) symmetry^14, 18, 19^. There are also a few algorithms for modeling asymmetric heteromeric complexes^20–22^. In addition, Multi-LZerD^23^ employs a genetic algorithm for heuristic conformational sampling, while the subsequent RL-MLZerD^24^ adopts reinforcement learning to enhance sampling efficiency. However, these methods can normally handle a limited number of chains.

Recently, the rapid advancement of deep learning has revolutionized the field of protein structure prediction^25–29^ and related applications^30–36^. AlphaFold2 (AF2)^25^, originally designed for monomers, has been adapted for complexes through several variants, including AF2-linker^37, 38^, FoldDock^39^ and AF2Complex^40^, which enable complex modelling without retraining. AlphaFold-Multimer (AFM)^41^, retrained specifically for protein complexes, demonstrates better predictive capabilities. However, AFM faces three major challenges when applied to large assemblies: marked accuracy degradation with increasing chain number and length^42^, limited conformational diversity for a given target, and strict memory constraints imposed by graphical processing units (GPUs). AFsample2^43^ enhances conformational diversity via random MSA column masking, but still does not address the GPU memory bottleneck. Running on an NVIDIA A100 GPU with 40 GB memory, AFM v2 can only handle sequences of up to 3300 amino acids. The recently released AlphaFold3 (AF3) server^44^ raises the input limit to 5000 residues. While AF3 supports local installation and execution now, the scale of complexes it can process remains constrained by GPU memory. On an 80 GB GPU, AF3 can predict up to approximately 5120 amino acids, which does not significantly exceed the capacity of AF3 server.

Due to the GPU memory limitation and accuracy reduction of AlphaFold in modeling large protein complexes, alternative strategies have emerged. Assembly-based methods first predict smaller subcomponents and subsequently assemble them into larger complexes. MoLPC, for example, employs a Monte Carlo Tree Search (MCTS) algorithm to assemble a full complex from dimers or trimers, achieving a success rate of approximately 30% with TM-score*>*0.8 on 175 large protein complexes^42^. CombFold applies a hierarchical and combinatorial assembly strategy, enabling broader exploration of potential inter-subunit interactions and yielding higher-quality predictions^45^. Nevertheless, these methods remain inherently dependent on the quality of AFM predictions, especially for homo-oligomers, where fewer transformations can be derived. Several recent methods aim to enhance the performance through additional constraints or learned prior knowledge. EvoDock integrates AFM-predicted models with symmetry constraints to model large complexes with cubic symmetry^46^. However, it depends on prior symmetry information. SymProFold^47^ is capable of predicting complete complexes with specific symmetries and unit cell parameters solely from input sequences. Yet, generalizing this approach to heteromeric or asymmetric complexes remains challenging. In addition, several methods have attempted to combine predicted models with other deep learning models^48, 49^, though none has yet demonstrated superior performance over CombFold.

To overcome current limitations, we develop HDOCK-Multimer (HDM), a fully automated framework that integrates docking and assembly strategies for accurate and efficient modeling of large protein complexes. On one hand, we introduce a genetic algorithm iteration into the combinatorial assembly process for better leveraging the correct pairwise interactions of predicted subcomplexes. On the other hand, we design an efficient multi-body docking algorithm based on our previously developed pairwise docking method, HDOCK^17^. In addition, given the success of previous docking methods in symmetry sampling, we also introduce a symmetric docking method to handle the symmetry of homo-oligomeric complexes. Such docking strategies can efficiently reduce reliance on subcomponent accuracy when AlphaFold yields incomplete or erroneous interfaces. Our HDM method is extensively validated on three benchmark datasets of large protein complexes, and achieves high top-1 success rates of up to 86% with TM-score*>*0.7, outperforming state-of-the-art assembly and deep learning-based methods.

## 2 Results

### 2.1 The workflow of HDOCK-Multimer

HDOCK-Multimer (HDM) takes the sequences of subunits as input and outputs the predicted complex structures. Figure 1a shows an overview of HDM, which is composed of three stages: (1) generation of monomeric and subcomponent structures using AF2/AFM, (2) determination of modeling strategies, and (3) structure modeling of the complex followed by scoring and clustering. In the first stage, we use AlphaFold2 (AF2) to predict all monomeric structures and AlphaFold-Multimer (AFM) to construct all subcomponent structures. Here, we define subcomponents as all possible trimeric subunit combinations, which can capture more possible cooperative interactions than pairwise combinations while avoiding expensive computational cost or GPU memory overflow of tetrameric combinations. AF2 outputs five models for each prediction, and only the top-ranked one is selected for subsequent tasks. For monomers, model selection is based on the average pLDDT values, whereas subcomponents are selected based on the model confidence scores (Equation 1). In the second stage, appropriate modeling approaches are determined based on the complex homology and pairwise docking results. Three modeling strategies are adopted in our study, including asymmetric docking, symmetric docking, and combinatorial assembly. Symmetric docking is used only for modeling homomeric complexes and supports a range of symmetry types, including cyclic (*C_n_*), dihedral (*D_n_*), tetrahedral (*T*), and octahedral (*O*) symmetries. Asymmetric docking is applicable to homomeric complexes as well as heteromeric complexes containing less than five unique chains. For complexes with five or more unique chains, only the assembly strategy is adopted because the docking accuracy drops sharply when the the number of chains increases. In the third stage, structure modeling and scoring are performed to build the complex structure of all subunits. We use our iterative knowledge-based scoring function^50^ to assess predicted multimeric systems, in order to score and rank the models generated by different prediction strategies. All complex models are clustered using a *Cα* root mean square deviation (RMSD) cutoff of 5 A° to reduce redundancy. Namely, if the RMSD between two models is within 5 A°, the one with the better score is kept. HDM outputs the top 10 models by default.

**Figure 1:**
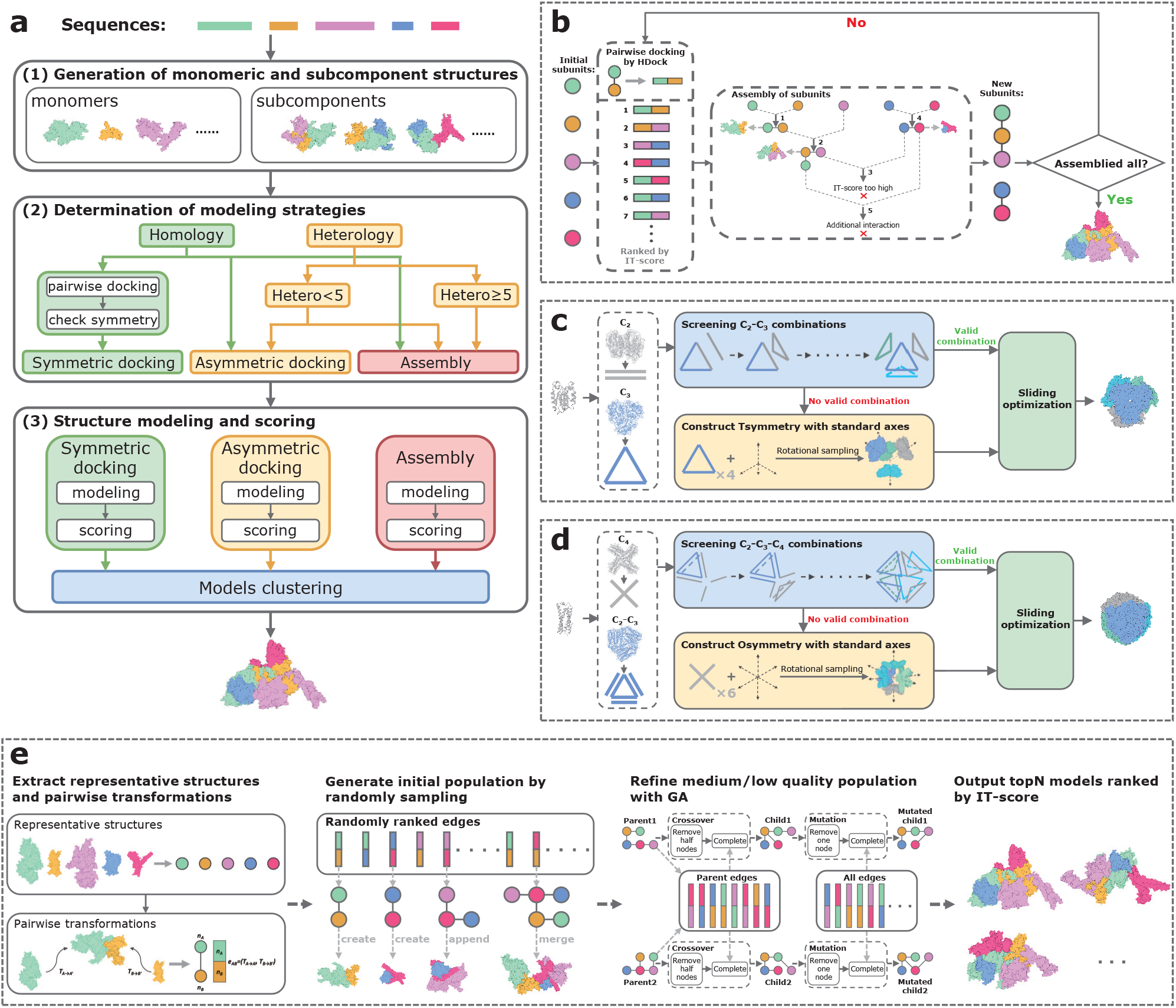
The framework of HDOCK-Multimer (HDM). **a**, The overview of HDM. The input is the sequences of subunits in the complex, and the output is the predicted complex structure. HDM integrates multiple modeling strategies, including asymmetric docking, symmetric docking, and assembly methods, to comprehensively explore the conformational space of protein complexes. **b**, The workflow of asymmetric docking algorithm. **c**, The workflow of tetrahedral docking algorithm. **d**, The workflow of octahedral docking algorithm. **e**, The workflow of combinatorial assembly algorithm.

### 2.2 Performance on Benchmark 1 (35 large heteromers)

HDM is first validated on Benchmark 1 of 35 large heteromeric complexes with 5 to 20 chains per complex. When evaluated by TM-score, HDM obtains a top-1 success rate of 83%, and accurately predicts 29 out of 35 complexes with TM-score *>* 0.7 (Fig. 2a), among which, 23 (or 66%) of the complexes yield high-quality predictions with TM-score *>* 0.8. Considering the top-10 predictions, the proportions of successful and high-quality models increase to 86% and 80%, respectively. In addition, HDM tends to perform better with the higher quality-weighted pairwise connectivity (QPC), giving a correlation coefficient of *r* = 0.70 (Supplementary Fig. 1a). HDM also shows a negative correlation with the increasing length of the complex (Supplementary Fig. 1b).

**Figure 2:**
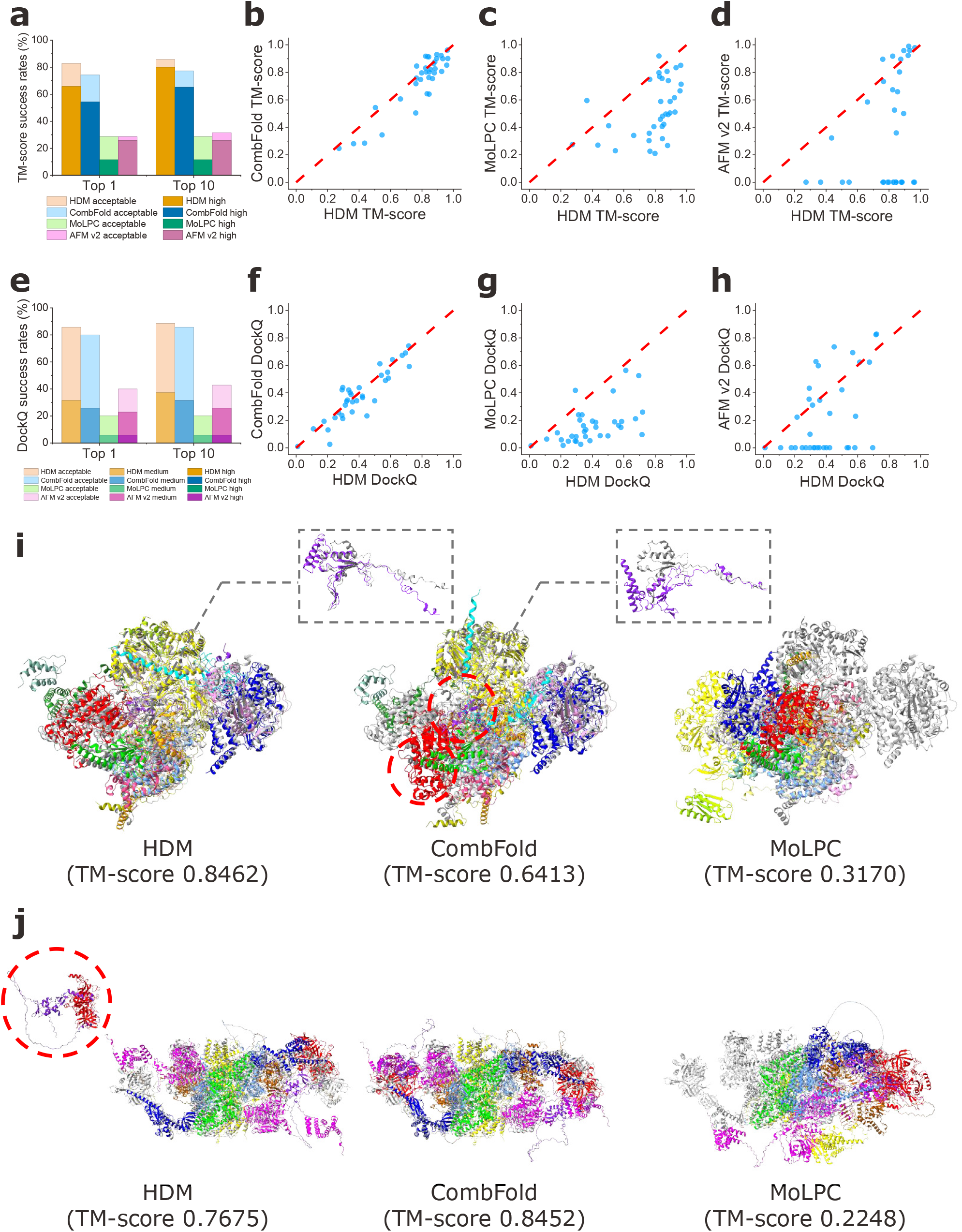
**Performance of HDOCK-Multimer (HDM) and other methods on Benchmark 1**. **a**, Top-*N* (*N* = 1, 10) success rates of HDM, CombFold, MoLPC and AFM v2, evaluated by TM-score. MoLPC produces only one prediction per target. **b**, TM-score comparison of the models predicted by CombFold and HDM. **c**, TM-score comparison of the models predicted by MoLPC and HDM. **d**, TM-score comparison of the models predicted by AFM v2 and HDM. **e**, Top-*N* (*N* = 1, 10) success rates of HDM, CombFold, MoLPC, and AFM v2, evaluated by DockQ. **f**, DockQ comparison of the models predicted by CombFold and HDM. **g**, DockQ comparison of the models predicted by MoLPC and HDM. **h**, DockQ comparison of the models predicted by AFM v2 and HDM. **i**, Polytomella Complex-I (PDB 7ARC): high-quality HDM model (left), inaccurate model for CombFold (middle) and MoLPC (right). The predicted models (colored by chain) are superimposed on the native structure (grey). Red dashed circles indicates those chains assembled into incorrect positions. The dashed boxes show enlarged views of wrongly assembled chain. **j**, EIF2B-eIF2 complex (PDB 6I3M): acceptable-quality HDM model (left), high-quality CombFold model (middle) and inaccurate MoLPC model (right). The predicted models (colored by chain) are superimposed onto the native structure (grey). Red dashed circles indicates those chains assembled into incorrect positions.

Compared with CombFold, MoLPC, and AFM v2 (AlphaFold Multimer version 2.2), HDM greatly outperforms the other methods (Fig. 2a). When evaluated by TM-score, HDM successfully predicts the acceptable and high-quality structures for 83% and 66% of the 35 cases, respectively, when the top-1 predictions are considered, which are substantially higher than 74% and 54% for CombFold, 29% and 11% for MoLPC, and 29% and 26% for AFM v2 (Fig. 2a). When considering the top-10 predictions, the success rates of HDM increase to 86% and 80% for acceptable and high-quality criteria, respectively, compared with 77% and 66% for CombFold, 29% and 11% for MoLPC, and 31% and 26% for AFM v2 (Fig. 2a).

The head-to-head comparison of the TM-scores between HDM and other methods reveals that HDM improves the performance over other methods in different ways (Fig. 2b-d). For HDM versus CombFold, HDM mainly improves the prediction on the cases when CombFold fails with TM-score *<* 0.7, whereas both methods perform comparably well on those cases where CombFold has a TM-score of over 0.7 (Fig. 2b). For HDM versus MoLPC, HDM greatly improves the TM-scores over MoLPC for most of the cases over the full range of TM-scores, and yields lower TM-scores than MoLPC for only two cases (Fig. 2c). For HDM versus AFM v2, HDM shows TM-scores comparable to AFM v2 on the cases where AFM v2 has a TM-score of 0.7 or better, while greatly improves the prediction on the remaining cases (Fig. 2d). It is especially noted that AFM v2 totally fails on quite a few cases due to the GPU memory limitations, while HDM is able to assemble the complexes with different degrees of accuracy (Fig. 2d).

Similar trends in performance improvement are observed when the DockQ metric is used (Fig. 2e). Overall, HDM outperforms the other methods, and achieves top-1 and top-10 success rates of 86% and 89% with acceptable accuracy, respectively, compared with 80% and 86% for CombFold, 20% and 20% for MoLPC, and 40% and 43% for AFM v2 (Fig. 2e). In addition, HDM yields higher DockQ values than the other methods for most of the cases (Fig. 2f-h). Interestingly, AFM v2 produces high-quality interfaces (DockQ *≥* 0.80) on a few targets like 7CK6 and 7Z15, outperforming HDM. This indicates that end-to-end approaches may still confer advantages in interface construction. Given the similar trends of HDM versus other methods in the success rates under TM-score and DockQ criteria, hereafter the TM-score metric will be used for evaluations by default to be consistent with that of previous studies ^42, 45^, unless otherwise specified.

Figure 2i,j shows a comparison of HDM with other methods for several examples of predicted complex structures. One example is the Polytomella Complex-I (PDB 7ARC), for which HDM gives a high-quality top-1 prediction with a TM-score of 0.85, whereas both CombFold and MoLPC fail with TM-scores of 0.64 and 0.32, respectively (Fig. 2i). For CombFold, the failure is primarily due to the misplacement of two subunits, as indicated by the red dashed circles and black dashed boxes in the Fig. 2i. Although HDM outperforms CombFold for most of the cases, there are a few cases in which CombFold achieves higher accuracy. For instance, CombFold assembles a high-quality model with a TM-score of 0.85 for the eIF2B-eIF2 complex (PDB 6I3M), while HDM yields a slightly worse but still successful prediction with a TM-score of 0.77 (Fig. 2j). The lower accuracy of HDM on this target mainly arises from an incorrect interface used during the assembly process, which leads to positional deviations of two subunits, as highlighted by the red dashed circles in Fig. 2j.

### 2.3 Performance on Benchmark 2 (172 large complexes)

HDM is further tested for its general applicability on Benchmark 2 of 172 large protein complexes including 114 homomers and 58 heteromers with 10 to 30 chains per complex. On this benchmark, HDM obtains a top-1 success rate of 59%, successfully modeling 102 out of 172 complexes with TM-score *>* 0.7, most of which (97 of 102) are of high quality with TM-score *>* 0.8 (Fig. 3a). When considering the top-10 predictions, the proportions of successful and high-quality predictions increase to 66% and 63%, respectively (Fig. 3a). In addition, HDM yields a comparable performance for both homomeric and heteromeric complexes and is not sensitive to the length of the complex (Supplementary Fig. 1d), suggesting the robustness of HDM.

**Figure 3:**
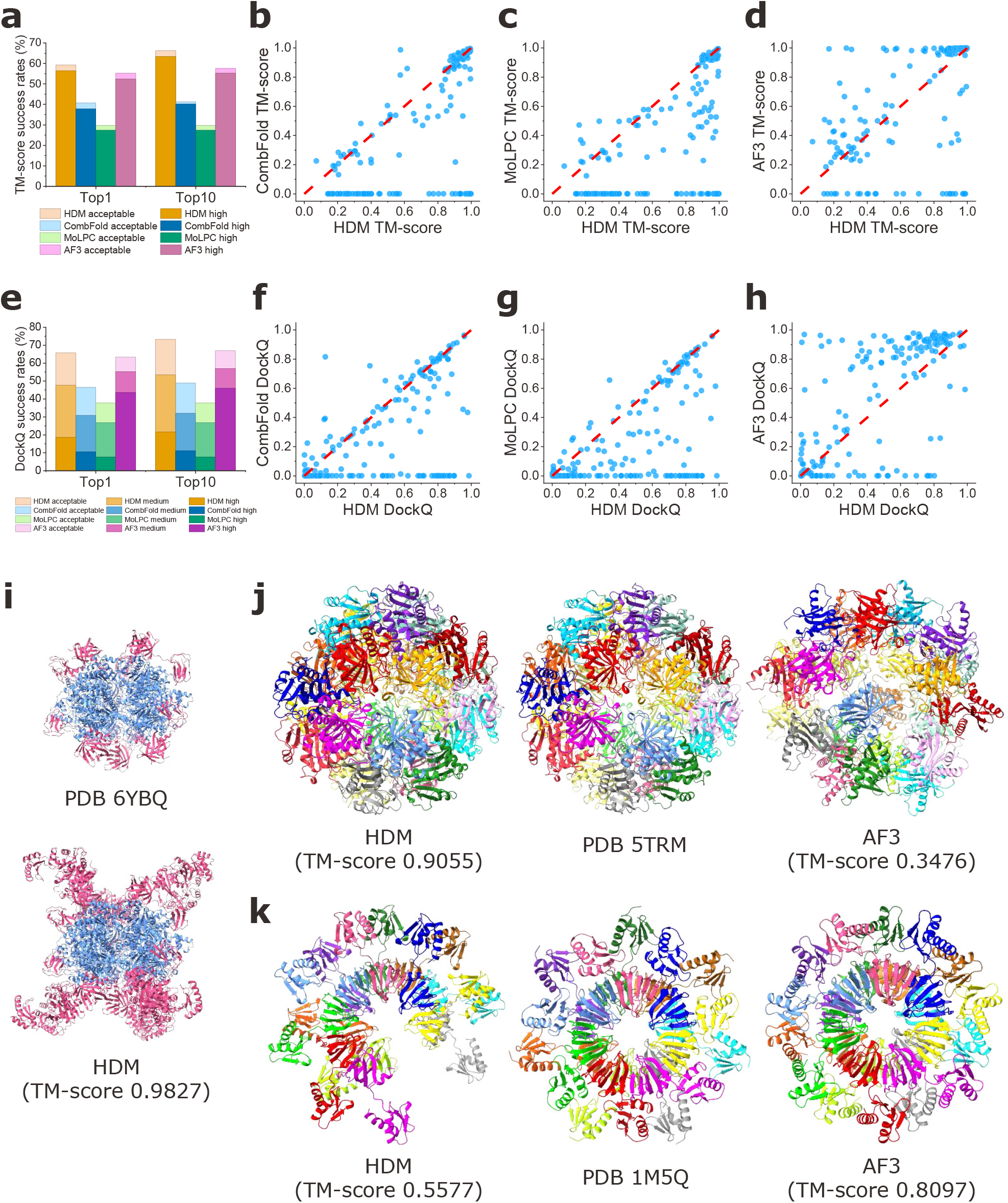
**Performance of HDOCK-Multimer (HDM) and other methods on Benchmark 2**. **a**, Top-*N* (*N* = 1, 10) success rates of HDM, CombFold, MoLPC and AF3, evaluated by TM-score. **b**, TM-score comparison of the models predicted by CombFold and HDM. **c**, TM-score comparison of the models predicted by MoLPC and HDM. **d**, TM-score comparison of the models predicted by AF3 and HDM. **e**, Top-*N* (*N* = 1, 10) success rates of HDM, CombFold, MoLPC, and AF3, evaluated by DockQ. **f**, DockQ comparison of the models predicted by CombFold and HDM. **g**, DockQ comparison of the models predicted by MoLPC and HDM. **h**, DockQ comparison of the models predicted by AF3 and HDM. **i**, Engineered glycolyl-CoA carboxylase (PDB 6YBQ): cryo-EM structure (top) and high-quality HDM model (bottom). HDM successfully predicted the structure of the unresolved regions of the *α* subunit in the cryo-EM structure, recovering a total of 2749 additional amino acids. **j**, Human GCN5 histone acetyltransferase domain (PDB 5TRM): high-quality HDM model (left), X-ray structure (middle), and inaccurate AF3 model (right). **k**, Augmented Sm-like archaeal protein from *Pyrobaculum aerophilum* (PDB 1M5Q): HDM-predicted model (left), X-ray structure (middle), and high-quality AF3 model (right).

Compared with CombFold, MoLPC, and AF3, HDM demonstrates the overall best prediction performance (Fig. 3a). Under the TM-score metric, HDM achieves the success rates of 59% and 66% for top-1 and top-10 predictions, respectively, compared with 41% and 41% for CombFold (Fig. 3a). MoLPC obtains a top-1 success rate of 30%. AF3 shows improved capability in complex modeling, yielding a top-1 success rate of 55%, with 90 out of 95 predictions classified as high-quality. When considering all five predictions, the success rate of AF3 increases to 58% (Fig. 3a). In addition, HDM also predicts more accurate models with higher TM-scores than the other methods for most of the cases (Fig. 3b-d). It is also noted that the other methods all fail on some complexes, while HDM can always predict the structure of the complexes with certain accuracies (Fig. 3b-d).

When assessing interface quality with DockQ, HDM still maintains the highest performance, giving top-1 and top-10 success rates of 66% and 73% with acceptable accuracy, respectively, compared with 47% and 49% for CombFold, 38% and 38% for MoLPC, and 63% and 67% for AF3 (Fig. 3e). In addition, HDM achieves higher DockQ scores than CombFold and MoLPC for most of the cases (Fig. 3f,g). Notably, although AF3 is able to model only a subset of the complexes (n = 145) due to input limitations, it produces substantially more high-quality predictions (Fig. 3h). The higher fraction of high-quality predictions highlights the advantage of end-to-end deep learning architectures in interface construction. However, it should be noted that the targets in Benchmark 2 are released before January 2022 and thus may overlap with the training data of AF3, which may contribute to its superior performance in generating high-quality predictions.

Compared with AF3, HDM is applicable to substantially larger protein complexes, and the incorporation of symmetric docking strategies further improves its ability to handle intricate symmetry. Here, we present two examples. Figure 3i illustrates the structure prediction of engineered glycolyl-CoA carboxylase (PDB 6YBQ) with 7062 amino acids, which exceeds the length limit of the AF3 server. For this complex, HDM not only built a high-quality prediction for the resolved regions of the cryo-EM structure (TM-score = 0.98), but also successfully modelled the previously unresolved portion of the *α* subunit, with an average pLDDT value of 91.06. Another example is the human GCN5 histone acetyltransferase domain (PDB 5TRM), the O-symmetric docking method of HDM produced a high-quality result (TM-score=0.91), whereas AF3 failed on this target (Fig. 3j). Nevertheless, the improved deep learning architecture of AF3, which allows efficient processing of both sequence and structural information, enables it to perform well on smaller complexes, accurately reconstructing the symmetry and complete structure. For instance, AF3 yielded a high-quality prediction (TM-score=0.81) for the Augmented Sm-like archaeal protein from *Pyrobaculum aerophilum* (PDB 1M5Q), although this complex (released in 2003) may be included in the training set of AF3, while HDM only predicted the partial complex with a TM-score of 0.56 due to the incorrect orientation predicted by the docking method (Fig. 3k).

### 2.4 Performance on Benchmark 3 (7 CASP15 targets)

To test HDM on real applications, we applied HDM on seven CASP15 targets including 2 homomeric complexes, 3 heteromeric complexes, and 2 proteins with a single chain longer than 3000 amino acids. For the single-chain targets, each chain was divided into domains according to IUPred3^51^. Disordered linker regions between domains were excluded from the prediction, and each domain was split equally to serve as input subunits. Using the automated pipeline, HDM produced acceptable-quality models for 6 of 7 targets (Fig. 4a-g), corresponding to a success rate of 86%.

**Figure 4:**
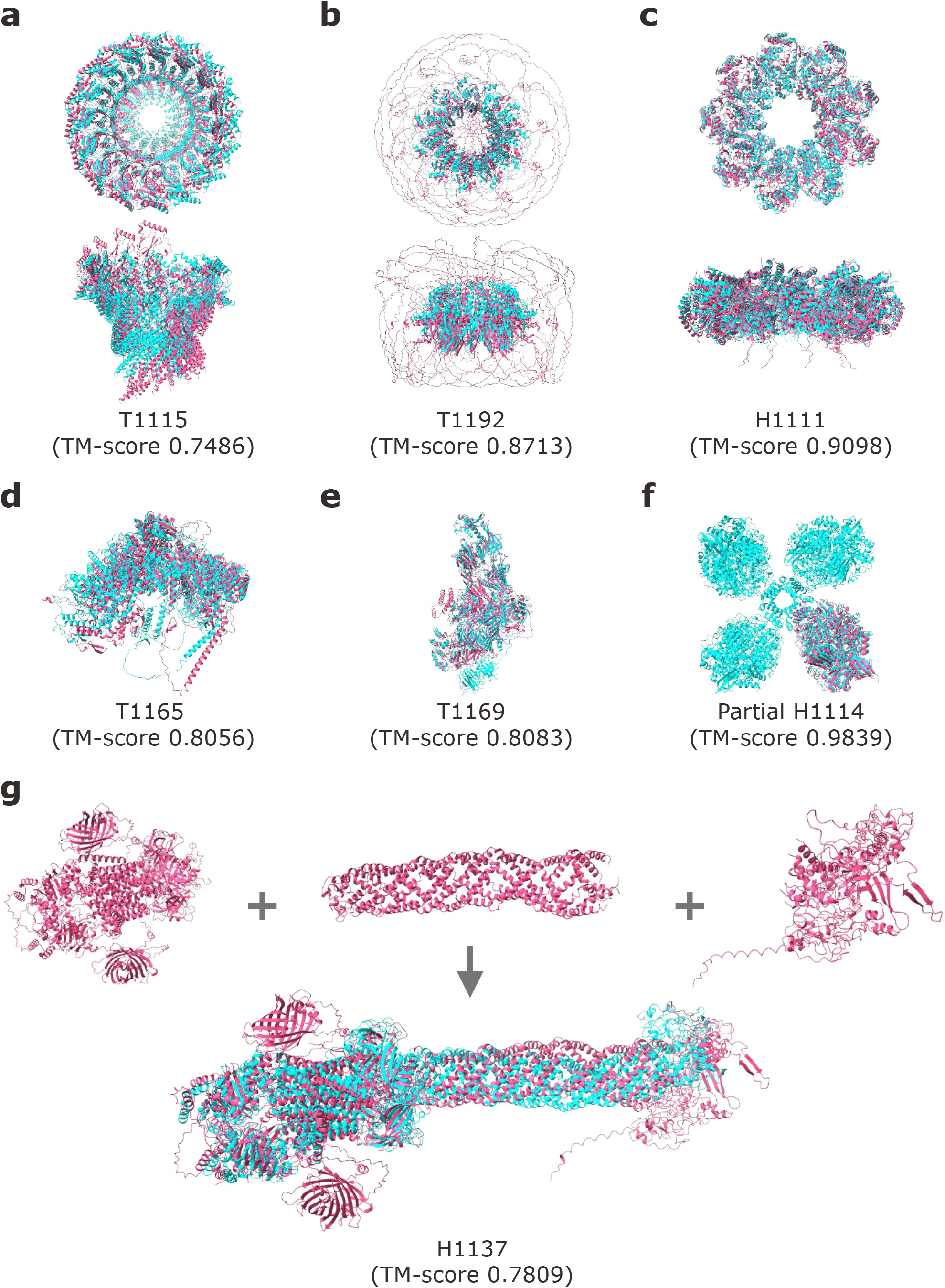
Predicted structures of HDOCK-Multimer (HDM) for 7 CASP15 targets of Benchmark 3. HDM predictions (pink) are superimposed on the experimental structure(cyan) or top-scoring CASP model if the experimental structure is not released. **a**, Acceptable HDM model of stomatin complex versus top-scoring CASP15 prediction. **b**, High-quality HDM model of RAD52 complex versus experimental structure (PDB 5XRZ). **c**, High-quality HDM model of YscY-YscX-LcrD complex versus experimental structure (PDB 7QIJ). **d**, High-quality HDM model of modified ligase Tom1 protein versus top-scoring CASP15 prediction. **e**, High-quality HDM model of native mosquito salivary gland surface protein 1 (SGS1) versus experimental structure (PDB 8FJP). **f**, High-quality HDM model of partial [NiFe]-hydrogenase Huc complex versus experimental structure (PDB 7UTD). **g**, Acceptable HDM model of MceG complex, obtained by connecting three seperately assembled domain groups, versus experimental structure (PDB 8FEF).

For H1111 (PDB 7QIJ), we removed the N-terminal domain of lcrD as it interfered with the remaining structure. This domain is also absent from the published experimental structure (Fig. 4c). For target H1114 (target 7UTD), while HDM did not recover the full complex, it yielded a high-quality partial subcomplex with a TM-score of 0.9839 (Fig. 4f). For target H1137 (PDB 8FEF), although the modeling with full chains as subunit input failed, we generated an acceptable model by manually dividing the chains into subunits (Fig. 4g). Specifically, AF2 was first employed to identify inter-chain interactions. Based on the predicted interactions, the full complex was partitioned into three groups, each comprising different chains or domain fragments (Supplementary Table 1). Adjacent domain fragments shared 20 overlapping amino acids, enabling structural alignment. HDM was then run independently on each group. The final structures were obtained by connecting models from each group via aligning the overlapping amino acids, followed by 1000 steps of MD relaxation.

Supplementary Table 2 shows a comparison of HDM with CombFold^45^, MoLPC^42^, AFM v2^41^, AF3^44^ and the top-ranked human and server groups in CASP15 competition. It can be seen from the table that HDM consistently outperforms MoLPC and AFM v2 which produces successful models for only three and one targets, respectively. HDM shows performance comparable to CombFold, sharing a success rate of 86%. HDM exceeds CombFold on H1111 but is inferior on H1137. Among the five targets within AF3’s input limit, HDM performs better on H1137 but worse on T1115. Notably, HDM achieves a higher success rate than AF3, owing to its ability of handling larger complexes. Overall, HDM demonstrates performance comparable to the leading competing methods.

### 2.5 Source of improvement over other methods

There are several differences in the assembly strategy of HDM compared with CombFold. First, the incorporation of a genetic algorithm (GA) optimization during the assembly process enables efficient exploration of the vast conformational space of large complexes while progressively filtering out the unfavorable binding orientations obtained in the initial sampling stage. By randomly swapping edges, only correctly oriented interfaces can be retained and participate in forming different structures without causing severe steric clashes. Second, the assembly process involves an IT-score scoring and re-ranking of final complexes after MD relaxation.

To investigate the advantages of the combinatorial assembly strategy in HDM over CombFold, we calculated the top-1 success rates of stepwise ablation models after each major step, including the basic “assembly” (i.e. the one similar to that of CombFold), “assembly+GA” (i.e. after GA optimization), “assembly+GA+MD” (i.e. after MD relaxation), and the full “assembly+GA+MD+ITscore” HDM baseline model (i.e. after IT-score scoring) on Benchmark 1. It is shown that the most basic “assembly” model performs better than MoLPC (Fig. 5a), suggesting that the assembly strategy of HDM is more efficient than the Monte Carlo search of MoLPC in exploring the binding space. Interestingly, the basic “assembly” model performs better than CombFold even they are pretty much the same (Fig. 5a), which could be attributed to the better pruning strategy in the assembly protocol of HDM than that of CombFold. In addition, the three components, including GA, MD, and IT-score, all contribute to the improvement of HDM over CombFold to some extent. Specifically, including “GA” improves the number of acceptable predictions over the basic “assembly” model. Although the MD relaxation does not improve the success rates over the “assembly+GA” model, it refines the assembled structures and reduces atomic clashes, which is necessary for the final step of IT-score scoring because IT-score is sensitive to clashes (Supplementary Fig. 2). Finally, the IT-score shows ability to better rank the relaxed predictions, leading to the improvement of the full “assembly+GA+MD+ITscore” model (i.e. HDM baseline) over the “assembly+GA+MD” model in both acceptable and high-quality predictions (Fig. 5a).

**Figure 5:**
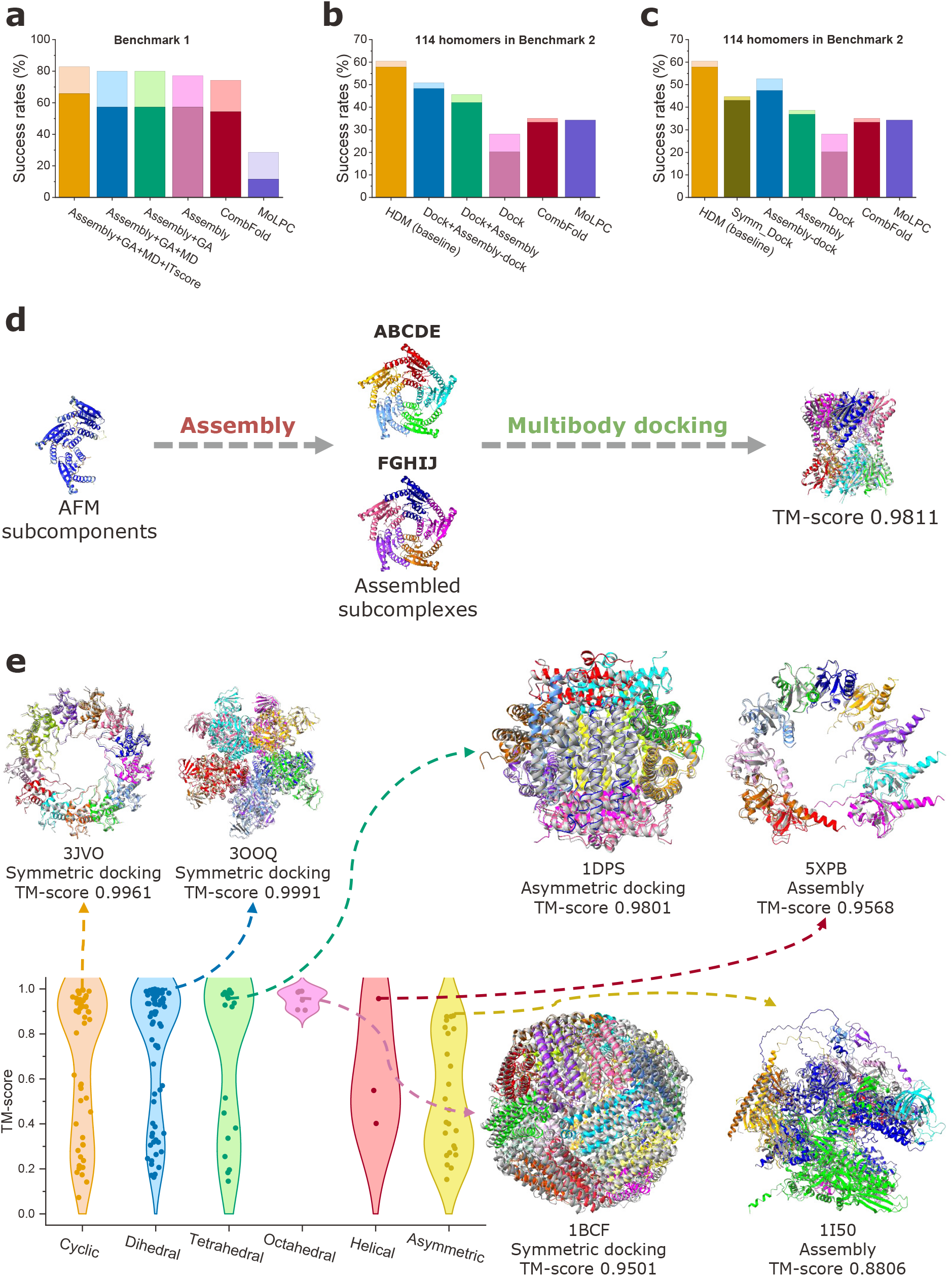
Advantages of HDOCK-Multimer (HDM) in integrating different strategies. **a**, Top-1 success rates of HDM assembly strategy, its stepwise ablation variants, CombFold and MoLPC on Benchmark 1, evaluated by TM-score. Starting from the full HDM assembly strategy (Assembly+GA+MD+ITscore), we sequentially remove IT-score ranking (Assembly+GA+MD), MD relaxation (Assembly+GA), and GA optimization (Assembly). **b**, Top-1 success rates of HDM (baseline), its stepwise ablation variants, CombFold and MoLPC on the 114 homomeric complexes of Benchmark 2, evaluated by TM-score. Starting from the full HDM framework, we sequentially remove symmetric docking (Dock+Assembly-dock), redocking of assembly-failed cases (Dock+Assembly), and assembly (Dock). **c**, Top-1 success rates of HDM (baseline), its ablation models, CombFold and MoLPC on the 114 homomeric complexes from Benchmark 2, evaluated by TM-score. The ablation models include multibody docking (Dock), symmetric docking (Symm Dock), assembly followed by multibody docking (Assembly-dock), and assembly only (Assembly). **d**, The “assembly followed by docking” trajectory for the lumazine synthase RibH2 from *Mesorhizobium loti* (PDB 2OBX). Intermediate subcomplexes and the final docked complex are colored by chain, with the final model superimposed onto the native structure (grey). **e**, TM-score distributions of HDM predictions and representative examples on Benchmark 2 grouped by symmetry type. Predicted models (colored by chain) are superimposed on the native structure (grey). The structures shown, with their corresponding symmetry types, prediction strategies, and TM-scores, are as follows: 3JVO (cyclic, symmetric docking, TM-score: 0.9961), 3OOQ (dihedral, symmetric docking, TM-score: 0.9991), 1DPS (tetrahedral, asymmetric docking, TM-score: 0.9801), 1BCF (octahedral, symmetric docking, TM-score: 0.9501), 5XPB (helical, assembly, TM-score: 0.9568), and 1I50 (asymmetric, assembly, TM-score: 0.8806).

In addition to the combinatorial assembly strategy (say “Assembly”), HDM also adopts docking strategies including asymmetric docking (say “Dock”) and symmetric docking (say “Symm Dock”). In addition, HDM also introduces a multi-body docking step to supplement the assembly process (say “Assembly-dock”) in case of assembly failure. To investigate the advantage of integrating multiple strategies in HDM, we calculated the top-1 success rates of stepwise ablation models after adding each strategy, including “Dock” only, “Dock+Assembly”, “Dock+Assembly-dock”, and “Dock+Assembly-dock+Symm Dock” (i.e. the HDM baseline model) on the 114 homomeric complexes of Benchmark 2. The performance improves progressively as additional strategies are incorporated, giving the top-1 acceptable and high-quality success rates of 28% and 20% for the ”Dock” model, 46% and 42% for the “Dock+Assembly” model, 51% and 48% for the “Dock+Assembly-dock” model, and 61% and 58% for the HDM baseline model (Fig. 5b). Among different strategies, the “Assembly” strategy obtains the most improvement, followed by “Symm Dock” and “Assembly-dock” (Fig. 5b). It is also noted from Figure 5b that both CombFold and MoLPC perform better than the basic “Dock” model, suggesting that both combinatorial assembly and Monte Carlo search strategies are more efficient than the docking process. However, after adding the “Assembly” strategy, the “Dock+Assembly” model outperforms both CombFold and MoLPC (Fig. 5b), indicating the advantage of integrating multiple modeling strategies.

To examine the contributions of different modeling strategies to HDM, we calculated the top-1 success rates of the ablation models using individual strategies on the 114 homomeric complexes of Benchmark 2. It can be seen from Figure 5c that the “Assembly-dock” protocol performs the best and gives the acceptable and high-quality predictions for 53% and 47% of the cases, followed by 45% and 43% for “Symm Dock”, 39% and 37% for “Assembly”, and 28% and 20% for “Dock”. From Figure 5c, one can also find that CombFold and MoLPC perform worse than all individual ablation models except the “Dock” model. Therefore, it can be understood that combining the advantages of multiple modeling strategies yields considerably higher success rates of HDM than CombFold and MoLPC (Fig. 5c).

It is noted that the “Assembly-dock” protocol is a supplementary step in addition to three basic modeling strategies for HDM. Such a process is particularly important for homomeric complexes, where AF2-predicted models often yield identical transformations between subunits, limiting the assembly to only partial structures of the complete complex. While these subcomplexes are of high accuracy, the absence of additional transformations prevents them from further extension. By introducing the docking procedure, HDM can utilize these high-quality subcomplexes to generate a full complex. For example, during the assembly of the lumazine synthase RibH2 from *Mesorhizobium loti* (PDB 2OBX), the assembly process could only produce two *C*_5_ symmetric subcomplexes using transformations extracted from AFM-predicted subcomponents (Fig. 5d). However, through further docking of these subcomplexes, HDM successfully generates a high-quality full complex structure (TM-score=0.98).

Furthermore, we analyzed the performances of HDM for different symmetry types on Benchmark 2 (Fig. 5e). It can be seen that for two most common symmetry types, cyclic and dihedral, as well as two higher-order symmetry types, T-symmetry and O-symmetry, HDM can successfully predict the majority of the complexes. Compared with symmetric complexes, asymmetric complexes tend to have lower TM-scores (Fig. 5e). Overall, symmetric complexes yield the top-1 and top-10 success rates 64% and 70%, compared with 33% and 42% for asymmetric complexes. The prediction is especially challenging on those partially symmetric targets. On one hand, relying on only one or two transformations from AF2-predicted models cannot accurately model asymmetric structures, and on the other hand, it is also difficult for the docking method to accurately distinguish different interaction modes between the same subunits. In general, integrating the predictions from different strategies can effectively enhance both the success rate and general applicability of HDM.

### 2.6 Stoichiometry prediction

Although stoichiometry information is needed by our method for structure prediction by default, our HDM assembly strategy has an ability to predict stoichiometry according to the confidence score defined in Equation 3 or the normalized IT-score of assembled structures. Here, the normalized IT-score is defined as the IT-score divided by the number of amino acids in the complex to eliminate size dependence. It is shown that the TM-score exhibits a modest Pearson correlation coefficient of 0.39 with the normalized IT-score on Benchmark 1 (Supplementary Fig. 3a). The receiver operating characteristic (ROC) analysis in identifying correction models yields an area under the curve (AUC) of 0.81 (Supplementary Fig. 3b), indicating the effectiveness of normalized IT-score in estimating model quality. We followed the similar stoichiometry prediction protocol used in CombFold^45^. Specifically, using the same AF2-predicted models as inputs, the assembly strategy enables rapid sampling of all potential stoichiometric combinations by enumerating different stoichiometries. The confidence scores or normalized IT-scores obtained after initial random sampling can be used to estimate the most likely stoichiometry or to significantly narrow down the range of possible stoichiometries. As such, our HDM method has an ability to build a complex without prior stoichiometry information.

Twenty representative complexes are selected from Benchmark 1 and 2 to evaluate the ability of HDM in predicting stoichiometry across diverse symmetry types, including four asymmetric heteromers, one helical assembly, five cyclic (*C_n_*) symmetric complexes, five dihedral (*D_n_*) symmetric complexes and five tetrahedral (*T*) symmetric complexes. For each target, we performed assembly using the same AF2-predicted models as inputs to traverse all potential stoichiometric combinations. The copy number of the repeated subunit was varied from 3 to 15, while maintaining the correct number of other subunits. As shown in Supplementary Table 3, the normalized IT-score achieves a high success rate of 70%, correctly predicting the stoichiometry for 14 out of 20 cases, compared with 45% and 9 cases for the confidence score (Supplementary Table 3). With the stoichiometry predicted by the normalized IT-score, HDM yields comparable TM-scores to that with explicit stoichiometry input, which is much better than the case using the predicted stoichiometry by the confidence score (Supplementary Fig. 3c,d). These results demonstrate the robustness of HDM on the cases with unknown stoichiometry.

One example for stoichiometry prediction is the bovine ATP synthase Fo domain (PDB 6ZBB), which contains eight copies of the mitochondrial ATP synthase F(0) subunit C1 forming a locally cyclic symmetry (Fig. 6a). We applied our assembly strategy to perform 500 rounds of random sampling across 13 candidate stoichiometries: 3–15 copies of subunit C1 and the correct number of copies for all the other subunits. Then, we compared the top-1 confidence scores among different stoichiometries. The complex with the correct stoichiometry of 8 copies for subunit C1 yields the highest confidence score (Fig. 6b), demonstrating the capability of using the confidence score to identify the correct stoichiometry during structure assembly with HDM. Similar phenomenon can also be observed in the normalized IT-score versus the number of subunit copies (Fig. 6c). Namely, the first minimum of the normalized IT-score happens at the correct stoichiometry of 8 copies, suggesting that the normalized IT-score can also be used to determine the correct stoichiometry.

**Figure 6:**
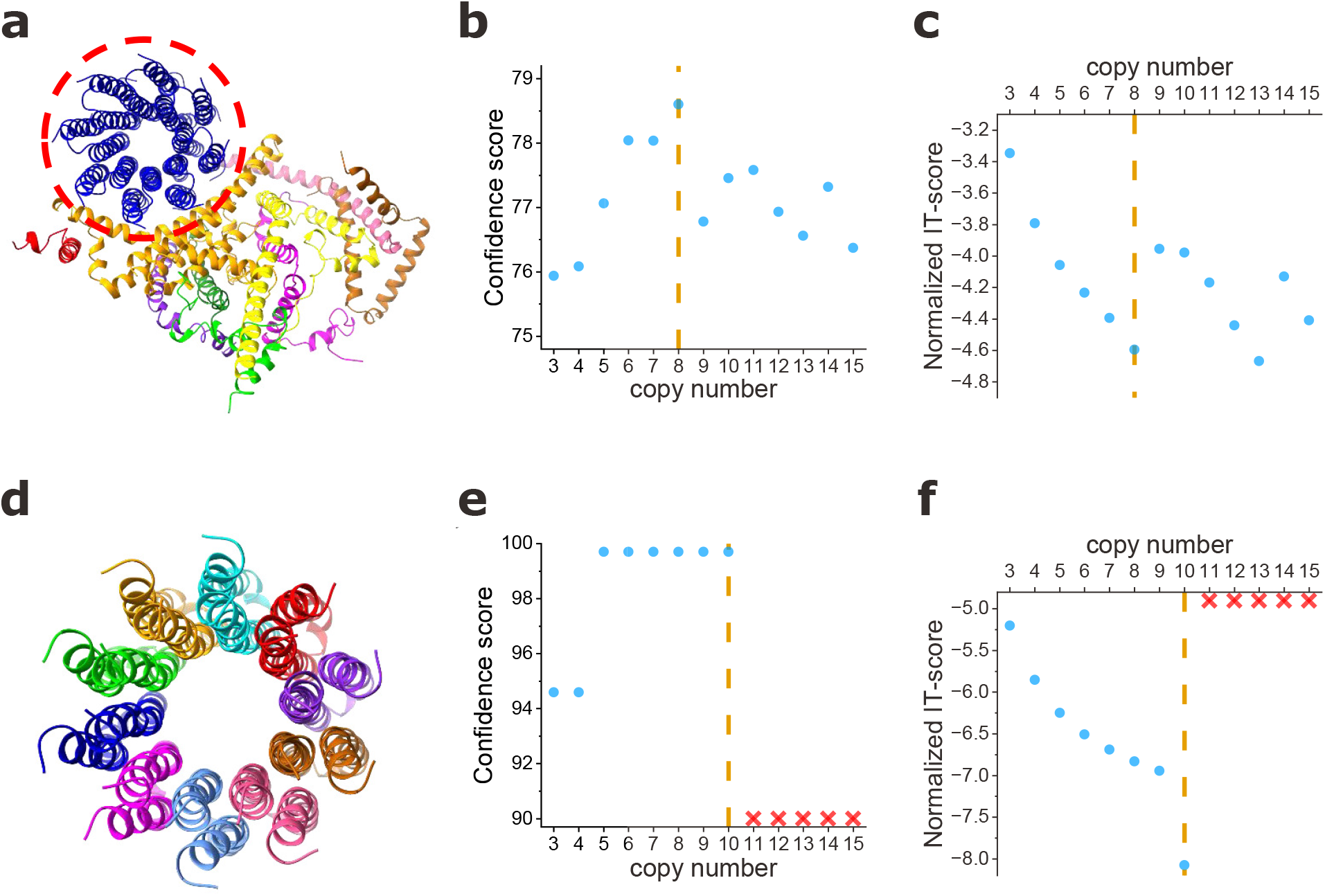
Stoichiometry prediction. **a**, Structure of the bovine ATP synthase Fo domain (PDB 6ZBB). A locally symmetric region formed by 8 copies of subunit C1 is marked with a red dashed circle. **b,c**, Top-1 predicted confidence score (**b**) and normalized IT-score (**c**) for the assembled structure as a function of the subunit copy number. The yellow vertical dashed line indicated the correct number of stoichiometry. **d**, Structure of the yeast F1Fo ATPase c10 ring with 10 copies of the subunit (PDB 4F4S). **e,f**, Top-1 predicted confidence score (**e**) and normalized IT-score (**f**) for the assembled structure as a function of the subunit copy number, where “*×*” stands for “failed to build a symmetric complex”. The yellow vertical dashed line indicated the correct number of stoichiometry.

Another example is the ring structure of the yeast mitochondrial ATP synthase that binds oligomycin (PDB 4F4S), consisting of 10 copies of mitochondrial ATP synthase subunit 9 (Fig. 6d). Using the same strategy, we conducted random sampling across 13 stoichiometries: 3–15 copies of subunit 9. Due to the symmetry of the trimer predicted by AFM, the top-1 confidence scores for complexes with 5 to 10 copies were all identical (Fig. 6e). However, for 11 or more copies, no complete complexes can be produced during sampling, likely due to steric clashes. These results suggest that, in addition to the confidence score, the ability to assemble the full complex can also be used to narrow down the plausible stoichiometry space. In contrast, the normalized IT-score exhibits superior discriminative capability. Among successfully assembled models with different stoichiometries, the one with correct stoichiometry of 10 copies yields the lowest normalized IT-score (Fig. 6f).

### 2.7 Training overlap and interface novelty analysis

Since HDM builds the complex structure from predicted subcomponents by AlphaFold-Multimer (AFM)^41^, we evaluated whether HDM performance might be influenced by interfaces already present in the training data of AFM or related models on Benchmark 1. Specifically, we searched the complexes in our benchmark against the PDB^52^ structures released before the AFM v2 training cut-off date (30 April 2018)^41^ using Foldseek-Multimer^53^. For each target in Benchmark 1, three types of query sets were generated from the complex structure: (i) all pairwise interfaces, (ii) all trimeric interfaces, and (iii) the full complex. Only the top-ranked hit was retained. We then calculated DockQ^54^ scores between the query and retrieved hit for assessing the interface similarity.

For each target, three interface similarity metrics were derived from the query sets: the mean DockQ value across all pairwise interfaces, the mean DockQ value across all trimeric interfaces, and the DockQ score of the full complex. We then analyzed the relationship between the HDM prediction accuracy and the interface similarity (Supplementary Fig. 4a-c). It is shown that there are only weak correlations between HDM TM-scores and interface similarities, with Pearson correlation coefficients of -0.11, -0.09 and 0.05 at the pairwise, trimeric, and full complex interface levels, respectively (Supplementary Fig. 4a-c). Notably, HDM is able to obtain high-quality predictions with TM-score *>* 0.8 for many cases with lower interface similarities of even below 0.1. These results indicate that the performance of HDM is not primarily driven by the interfaces that already have been seen by AFM, supporting the capability of HDM to assemble novel complexes.

## 3 Discussion

We have proposed a hybrid modeling framework for structure prediction of large protein complexes by integrating ab initio docking and combinatorial assembly from predicted subcomponent structures. Compared with state-of-the-art methods, HDM demonstrates superior or comparable performances on three benchmark datasets, including 35 heteromeric complexes comprising 5 to 20 chains and 1,300– 8,000 amino acids per complex, 172 general complexes with 10 to 30 chains and 500-10,000 amino acids per complex, and seven CASP15 targets. Analysis of different modeling strategies reveals that incorporating docking methods not only increases the success rate of the assembly process, but also reduces the dependence of the HDM results on AFM prediction quality, thereby enhancing the general applicability of our HDM method. HDM can not only be used for structure prediction of multi-chain complexes by ab initio docking, but also serve as a powerful complement to AF2/AF3, overcoming GPU memory limitations for the prediction of larger protein complexes. Moreover, it can also recover the unresolved regions that are missing in some structures by experiment methods.

Despite the present success of our HDM method, there are still room to improve in the future work. First, our symmetric docking method only covers a subset of symmetries. Other symmetry types, such as helical symmetry, are not yet supported and rely on the transformations predicted by AFM. Second, some model ranking failures indicate that simply summing up pairwise interaction scores is insufficient to capture the intricate multi-body interactions in large complexes. Current methods largely rely on pairwise interface metrics and may not capture higher-order cooperativity or collective effects in large complexes. Therefore, there is an urgent need to develop a true multi-body scoring function. Third, the performance of the multi-body docking algorithm still has room for improvement, especially for heteromeric complexes. Incorporating AF2-or AF3-predicted interface information during early docking stages may provide useful guidance. Fourth, the current method still relies on known stoichiometric information, although we have demonstrated the ability of our assembly method in predicting or narrowing down the stoichiometry space. Enhancing this capability will much reduce the computational cost and improve the accuracy for modeling complexes with unknown stoichiometry. Fifth, the AFM-derived confidence score is currently adopted to select the subcomponent structures for assembly and docking. Other metrics like Local Interaction Score (LIS)^55^ and predicted DockQ score (pDockQ)^56^, which demonstrate advantages over the confidence score, could also be used to selecting the best subcomponent model. Sixth, for convenience, we used RMSD to cluster sampled models in the current HDM protocol. Other metrics like fraction of common contacts or orientational RMSD could also be explored to remove the redundant complexes.

Although HDM shows an ability to predict stoichiometry, it should be emphasized that none of these tools (HDM, CombFold, etc.) provide truly absolute stoichiometry predictions, as this ultimately relies on the structure quality of predicted subcomponents. Relative stoichiometries may be inferred from experimental techniques such as mass spectrometry, but current computational approaches remain limited in this regard. Therefore, integrating experimental data, such as crosslinking mass spectrometry^57^, fluorescence resonance energy transfer (FRET)^58^ and cryo-EM^59^, into the prediction process, may further improve the modeling accuracy. Currently, HDM is able to utilize the cross-linking information, and demonstrates the potential to improve the modeling accuracy after incorporating provided cross-linking constraints on target 7EGF (Supplementary Fig. 5). The future study will focus on incorporating other experimental data. Examination of our results reveals that HDM has an ability to model antibody-antigen complexes. Among four antibody-antigen targets in Benchmarks 1 and 2, HDM achieves successful predictions on two targets (6SSI and 4RDQ), but fails on the other two (2VYR and 6SSI) (Supplementary Table 4). A more extensive evaluation on antibody-antigen complexes is needed to demonstrate the robustness of HDM on such targets.

## 4 Methods

### 4.1 Benchmark datasets

Three benchmark data sets are used to validate our HDM method. Benchmark 1 is taken from the study of CombFold^45^, consisting of 35 heteromeric complexes with 5 to 20 chains and 1,300 to 8,000 amino acids per complex (Supplementary Fig. 6a). All complexes in Benchmark 1 were released after April 2018, and thus not included in the training set of AFM v2^41^. Benchmark 1 is used to assess the performance of HDM on large heteromeric complexes. On this dataset, we compared HDM against CombFold, MoLPC^42^, and AFM v2. The second benchmark, named Benchmark 2, is derived from the study of MoLPC. After removing the same complexes in Benchmark 1, the resulting set contains 172 complexes with size ranging from 500 to 10000 amino acids and 10 to 30 chains per complex (Supplementary Fig. 6b). Among these, 114 are homomeric and 58 are heteromeric assemblies. In terms of symmetry, Benchmark 2 includes 48 cyclic (*C_n_*), 70 dihedral (*D_n_*), 21 tetrahedral (*T*), 6 octahedral (*O*), 3 helical (*H*) symmetric complexes, as well as 24 asymmetric complexes. All complexes in Benchmark 2 were released before January 2022. This dataset enables a broader evaluation of HDM, extending the assessment to larger complexes and more intricate symmetries. On benchmark 2, HDM was compared with CombFold, MoLPC, and the latest end-to-end deep-learning method, AF3^44^. Benchmark 3 consists of seven CASP15^60^ targets with more than 3,000 amino acids, including 2 homomers, 3 hetermomers and 2 single-chain proteins. Benchmark 3 represents a more challenging evaluation category in real applications. In addition to the methods mentioned previously, HDM was further compared with the top-ranking competitors from CASP15 on this benchmark.

In this study, the smallest component used for docking or assembly is called subunit that can be a full chain or part of a long chain. If a case exceeds the GPU memory limitation of AlphaFold, its chains will be broken into subunits. Across three benchmark datasets, nearly all targets use full-length chains as subunits. For the yeast Swi/Snf complex in a nucleosome-free state (PDB 7C4J) in Benchmark 2, the two long chains are divided into two subunits based on sequence length: the first 1000 amino acids formed the first subunit, and the remaining residues formed the second. This division ensures that the resulting subcomponents remain within the length constraints of AFM (AF2). For two long single-chain targets in Benchmark 3 (T1165 and T1169), each chain was partitioned into domains according to IUPred3^51^. Predicted disordered linker regions between domains were excluded from the prediction, and each domain was subsequently split into equally sized segments to serve as input subunits. Notably, during the docking stage, the full-length chains were still used as subunits. As such, our HDM method can predict the structure of large protein complexes of any sizes.

### 4.2 AlphaFold structure prediction

Here, AlphaFold is used to predict the structures of monomers or trimeric subcomponents. The choice of trimeric subcomponents is based on a balance between accuracy and speed because dimeric subcomponents will result in reduced modeling accuracy (Supplementary Fig. 7) while predicting all tetrameric subcomponents is computationally expensive or even prohibitive. Specifically, we first conduct a sequence search in the AlphaFold Protein Structure Database (AFDB)^61^ to check whether predicted monomer structures are available. If a sequence-matching hit is retrieved, we retain the structure with the highest average pLDDT value. If there is no available structure in the AFDB, structure prediction of the monomer is performed using AF2, and the model with the highest pLDDT is selected for subsequent docking. Next, all possible trimeric subunit combinations are generated using AF2. We have verified that replacing the software used to predict subcomponent structures, e.g. from AF2 to AF3, does not significantly affect HDM performance (Supplementary Fig. 8). Only the highest-confidence subcomponent structures are retained for the subsequent assembly process. The model confidence of AF2-predicted multimer is defined as:

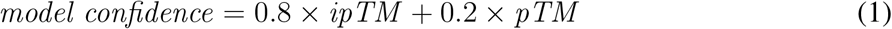

For comparison purpose, AF2 and AF3 are also used to predict the structures of the complexes in Benchmarks 1 and 2, respectively.

It should be noted that the disordered terminal regions of AF2-predicted monomers may interfere with the docking process. These regions may cause steric clashes, leading to correct orientations being assigned worse scores. Therefore, we apply a preprocessing step to remove disordered terminal residues from predicted monomer structures. First, we calculate the average pLDDT value for each chain. For chains with an average pLDDT *≥* 80, we remove terminal residues with pLDDT *<* 70. For chains with an average pLDDT between 50 and 80, secondary structure is predicted by STRIDE^62^, and terminal residues are removed if their plDTT *<* 50 or do not belong to a helix or sheet. For chains with an average pLDDT *<* 50, the full-length structure is retained without any trimming. After the complex structure is generated, such disordered terminal regions can be added back by approaches like homology modeling, MD simulations or structure alignment (Supplementary Fig. 9).

### 4.3 Modeling strategies

Three modeling strategies, asymmetric docking, symmetric docking, and combinatorial assembly, are adopted to build the full complex structure of all subunits, one of which will be chosen at a time by considering several factors. First, all input sequences are compared to determine whether the complex is homomeric or heteromeric. For homomeric complexes and heteromeric complexes with fewer than five unique chains, asymmetric docking and assembly strategies are selected by default. For complexes composed of five or more unique chains, only the assembly strategy is employed. For the cases with structural symmetry, symmetric docking is chosen to build their homomeric complex structures (Fig. 1a).

As the exact symmetry type may be unknown in realistic applications, HDM does not require explicit specification of symmetry information. If the symmetry is not provided, possible symmetry combinations for symmetric docking strategy, including cyclic, dihedral, and cubic symmetries, are inferred from the number of chains and pairwise docking results. To identify possible symmetric configurations, the total number of chains, *N*, is factorized as *N* = *n*_1_ *× n*_2_, where each factor pair (*n*_1_, *n*_2_) corresponds to two global cyclic symmetries, *C_n_*_1_ and *C_n_*_2_ . Specifically, the pair (1, *N*) corresponds to *C_N_* symmetry; if *n*_1_ = 2, both *C_n_*_2_ and *D_n_*_2_ symmetries are considered. For *N* = 12 or 24, tetrahedral (*T*) or octahedral (*O*) symmetries are also included.

Next, we perform pairwise docking of subunits using HDOCK and detect the rotational symmetry of docking results to further constrain the symmetric search space. The top 10 transformations from docking results are iteratively applied *m* times (*m* = 1, 2*, . . ., N −* 1) to generate the structures containing *n* chains (*n* = 2, 3*, . . ., N*). For each *n* that divides *N*, steric clashes are checked, defined as the total number of backbone atom pairs within 2 A° across all chain pairs. Structures with clashes below the following threshold are retained:

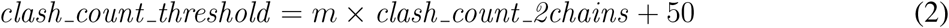

where *clash count 2chains* represents the number of clashes in the dimer (*m* = 1).

For filtered candidates, potential *C_n_* symmetry is further assessed. The centroid of each chain is computed to identify potential *C_n_* symmetry axis. Each chain is then rotated by 2*π/n* around the axis, and the *Cα* RMSD between the rotated and adjacent chains is calculated. Structures with RMSD *≤* 2 A° are considered *C_n_*-symmetric. If multiple *C_n_* symmetries are detected, the highest-order one is retained, denoted as *C_n_* (*pair*). To validate the detected symmetry, a specialized *C_n_*-symmetric docking is performed using CHDOCK. Among the top-10 CHDOCK results, the structure that can be superimposed onto *C_n_* (*pair*) within 2 A° *Cα* RMSD are selected, denoted as *C_n_* (*chdock*). The lowest-energy *C_n_* (*chdock*) model is retained for asymmetric docking, while the factor pair (*n, N/n*) determines the symmetric configurations. If no valid *C_n_* symmetry is identified, the monomer is used for asymmetric docking directly, and all possible symmetry types will be explored. Taking *N* = 12 as an example, if *C*_2_ symmetry is detected, *C*_2_, *C*_6_, *D*_6_, and *T* symmetries are all tried during the modeling stage. If *C*_3_ is detected, *C*_3_, *C*_4_, and *T* symmetries are sampled. No symmetry detection requires exploring all seven symmetries (*C*_2_*, C*_3_*, C*_4_*, C*_6_*, C*_12_*, D*_6_*, T*).

### 4.4 Asymmetric docking

The asymmetric docking method is developed based on our rigid-body protein-protein pairwise docking method, HDOCK^17^. The docking procedure of HDOCK is based on a fast Fourier transformation (FFT)-based search strategy. Specifically, the receptor is fixed and the ligand is rotated by an interval of Euler angles (*Δϕ*, *Δθ*, *Δψ*). During the rotational sampling, an angle interval of 15*^◦^* is adopted, resulting in 4392 orientations. At each rotational orientation, both proteins are projected onto a three-dimensional (3D) grid of N *×* N *×* N grid points with 1.2 A° grid spacing for translational search. The grid values incorporate contributions from both nearest neighboring points and long-range interactions in the form of *e*^−*r*2^. The calculation of shape complementarity is accelerated using 3D FFT^63^, and the top-10 translations with the best complementarity scores for each rotation are further refined using our iterative scoring function, IT-score^50^. The best-scored docking pose is retained for each rotation, yielding a total of 4392 binding modes.

We propose a multi-body docking strategy based on HDOCK for predicting protein assembly structures without symmetry constraints (Fig. 1b). Each chain is treated as a node *n_i_* (*i ∈* [1*, N*]), and pairwise docking interactions are considered as edges *e_ij_* (*i, j ∈* [1*, N*]). Thus, the complex modeling problem becomes a minimum spanning tree (MST) construction problem. To handle spatial constraints, our method integrates the principles from Prim’s^64^ and Kruskal’s algorithm^65^, and iteratively updates edge scores based on docking predictions to guide assembly.

HDOCK first performs pairwise docking for all input chains, with only the best-scored transformation retained as the initial edge for each pair. These edges are sorted in ascending order of IT-score, and the MST construction starts from the best-scored edge. When adding a new node *n_j_* to an existing partial spanning tree *T_k_* through edge *e_ij_*, both docking score screening and interaction interface screening are applied. The IT-score of *e_ij_* cannot exceed half of the lowest edge score in *T_k_* (denoted as *min score_k_*), i.e., *IT score*(*e_ij_*) *≤ min score_k_/*2; otherwise, the expansion of *T_k_* is terminated.

In addition, if more than 30 backbone atoms of *n_j_* are within 10 A° of any chain in *T_k_* except *n_i_*, the expansion is discarded to avoid undesired interfaces. In most cases, the entire complex cannot be built in the first iteration, resulting in several subcomponents, such as monomers, dimers, and trimers. They will be treated as new nodes, and pairwise docking is repeated to update the interaction graph. The docking and screening cycle is iterated until the full assembly is achieved. For homomeric complexes or complexes containing homologous chains, the asymmetric docking process can be significantly simplified, as identical chains require only a single docking run. This enables simultaneous incorporation of multiple chains, reducing both computational time and iteration cycles.

### 4.5 Symmetric docking

For symmetric docking, we used our previously developed *C_n_* and *D_n_* symmetric docking methods, CHDOCK^66^ and DHDOCK^19^, both implemented as rigid-body docking approaches. CHDOCK applies symmetry constraints during rotational sampling by setting the z-axis as the rotational symmetry axis. The sampling range in the rotational space is reduced to (*ϕ* = 0, *θ ∈* [0*, π/*2], *ψ ∈* (0, 2*π*]), with an interval of 10*^◦^*, resulting in 360 orientations. The translational sampling and scoring procedures are similar to those in HDOCK, ultimately producing 360 *C_n_* symmetric binding modes. DHDOCK performs additional *C*_2_ symmetric sampling using a cyclic axis perpendicular to the *C_n_* symmetry axis (z-axis) on the basis of *C_n_* symmetric models generated by CHDOCK.

### 4.5.1 *C_n_*/*D_n_* symmetric docking

To construct a *C_n_*_1_ symmetric structure with *N* chains, we adopt a stepwise cyclic symmetry sampling strategy, yielding a *C_n_*_2_ -*C_n_*_1_ symmetric mode, where *N* = *n*_1_ *×n*_2_. Here, *C_n_*_2_ and *C_n_*_1_ represent local and global symmetries, respectively. In the local symmetry sampling stage, CHDOCK performs *C_n_*_2_ symmetric docking on the monomer, and the top-10 models are retained. Global sampling applies *C_n_*_1_ symmetric docking to each *C_n_*_2_ mode, generating 100 candidates in total. These models are reranked by IT-score and clustered, and the 10 best-scored structures are selected as the final output. For *C_N_* symmetry, only a single round of cyclic symmetry sampling is needed. *D_n_*_1_ symmetric complexes are modelled using DHDOCK via a similar workflow, yielding the top-10 structures. The main difference between *D_n_*_1_ and *C_n_*_2_ -*C_n_*_1_ modes lies in the orthogonal orientation of the cyclic axes^67^ in two sampling stages.

#### 4.5.2 ***T*** -symmetric docking

Tetrahedral (*T*) symmetric structures, characterized by both *C*_2_ and *C*_3_ symmetries, consist of 12 chains for the simplest homomeric case. Therefore, for homomeric complexes with 12 chains, additional T-symmetric docking is performed. First, *C*_2_ and *C*_3_ symmetric docking are applied to generate candidate local symmetric modes. Then, *C*_2_-*C*_3_ combinations, which are capable of forming T-symmetry, are identified through the following process: (1) superimposing the first chain of a dimer onto a trimer, (2) expanding a new trimer using the second chain of the dimer as an anchor, and (3) verifying whether the newly expanded chains form *C*_2_ symmetry. Valid combinations undergo sliding optimization, where each trimer is translated along its cyclic axis to minimize steric clashes. For each combination, the best-scored structure is retained. The top-10 structures are selected as the final output. If no valid combination is found, T-symmetric systems are constructed using trimeric structures and standard tetrahedral axes. Each candidate trimer is replicated three times, with their *C*_3_ axes aligned to those of a standard tetrahedron. These trimers are then rotated around their axes in steps of 15*^◦^* until *C*_2_ symmetries are formed between different chain pairs, followed by sliding optimization to obtain the final outputs (Fig. 1c).

#### 4.5.3 ***O***-symmetric docking

For homomeric complexes composed of 24 chains, we perform O-symmetric docking. *C*_2_-*C*_3_ and *C*_4_ symmetric docking are first conducted to generate candidate local symmetric models. Next, combinations capable of forming O-symmetry are identified through a similar process: (1) superimposing two chains of a tetramer onto a hexamer, (2) expanding a new hexamer using the remaining two chains as anchors, (3) verifying *C*_4_ symmetries of the non-anchor chains in the new hexamers. Valid hexamer sets also undergo sliding optimization to obtain the top 10 structures. If there is no valid *C*_2_-*C*_3_-*C*_4_ combination, O-symmetric systems are constructed using tetramer structures and standard octahedral axes (Fig. 1d).

### 4.6 Genetic algorithm-based combinatorial assembly

The assembly workflow of HDM is illustrated in Fig. 1e, which consists of three stages: (1) extracting representative subunits and pairwise transformations, (2) initial random sampling, and (3) refinement via genetic algorithm (GA) iteration. The third stage is only applied when the initial sampled structures exhibit suboptimal confidence. Final models can be relaxed optionally to reduce steric clashes.

#### 4.6.1 Extracting representative subunits and pairwise transformations

First, all subunits are extracted from AFM-predicted subcomponents and ranked by their average pLDDT scores. The highest-ranking structures are selected as the representative subunits. Then, pair-wise interactions are identified between subunit pairs if any *Cα*-*Cα* distance is within 8 A° . Once an interaction is detected, transformations from the representative subunits to the corresponding structures are calculated. Considering possible structural variations in identical subunits from different models, alignment is performed by MMalign^68^ to obtain more precise transformations. To improve alignment, residues with pLDDT *<* 80 are removed from representative subunits with median pLDDT *>* 90. Each transformation is scored by the average pLDDT values of interface resides. Local symmetry often exists in the complexes with multiple identical chains. Therefore, for complexes with five or more identical subunits, transformation scores extracted from homomeric subcomponents are rescaled using *S* = *S* + *S ×* (1 *− S/*100), where *S* is the original score.

#### 4.6.2 Initial sampling stage

This stage includes 500 independent random sampling rounds. In each round, all transformations are randomly shuffled and traversed sequentially. Three assembly operations are applied to construct new structures:

- *Create*: assemble two unpaired subunits.
- *Append*: incorporate an unpaired subunit into an existing subcomplex.
- *Merge*: combine two different subcomplexes into a larger complex.

To avoid steric clashes from inaccurate transformations, clash examination is performed for the backbone atoms of residues with pLDDT *>* 80 in each operation. Clashes are defined as backbone atoms whose distance are within 2 A° across different subunits. If the clash ratio exceeds a predefined threshold, the operation is discarded. Thresholds vary by operations to account for the effects of subcomplex size: 2% of the shorter subunit/subcomplex length for *Create* and *Merge*, and 5% of the unpaired subunit length for *Append*. For subunits from the same chain, additional sequence connectivity is checked. The operation is discarded if the distance between consecutive residues from two subunits exceeds the number of linker residues multiplied by 4 A° .

Successful assembly requires all chains to be included into a single complex. For fully assembled complexes, the confidence score is calculated as a weighted transformation score^45^, and defined by the following expression:

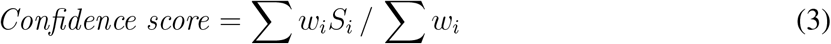

where *S_i_*represents the transformation score, and the weight *w_i_* is the number of residues in the shorter subunit or subcomplex in each assembly operation.

Each failed sampling yields a set of partial complexes, which is assigned a confidence score based on its largest subcomplex. After 500 iterations, fully assembled complexes are ranked by their confidence scores and clustered. The highest-confidence structure in each cluster is retained as the representative. The top 100 representatives are selected to form the initial population. The population quality is represented by the lowest confidence score of the top 20 structures, denoted as *S_PQ_*, and classified into three categories: high quality (*S_PQ_ ≥* 85.0), medium quality (80.0 *≤ S_PQ_ <* 85.0), and low quality (*S_PQ_ <* 80.0). Medium-and low-quality populations proceed to the GA optimization stage. High-quality populations or those with fewer than 10 clusters are re-ranked by IT-score and clustered, with the top representatives selected for output. If all 500 iterations fail to build the full complex, the set of structures containing the largest and highest-confidence subcomplex is retained. Each chain in the set is filtered using the same criteria as in monomer preprocessing, and the full complex is subsequently obtained using the multi-body docking algorithm.

#### 4.6.3 GA optimization stage

We adopt a genetic algorithm (GA) to optimize medium-and low-quality populations. In this work, the population size *M* is set to 100, and the mutation rate is set to 0.1. Two GA operations, crossover and mutation, are used to generate new structures. Crossover randomly selects two parent structures, removes up to half of the subunits from each, and reconstructs the complex by reapplying transformations from parents. Each crossover operation can yield up to two new candidates. Mutation randomly removes a subunit from a newly generated structure and rebuilds the complex using the transformations from the full set. Both operations follow the same assembly process as that for the initial sampling stage. The clash threshold for *Append* operation is adjusted to 2% in mutation to reduce the risk of erroneous operation.

Each GA generation performs 2*M* (i.e. 200 here) operations. All structures are ranked by confidence score and then clustered. The top-*M* representatives are retained for the next generation. Unlike previous GA-based approaches that use fixed procedures, we adopt a dynamic termination strategy that stops the iteration once the population quality improves by one level (i.e., medium-quality populations continue until *S_PQ_ ≥* 85.0; low-quality until *S_PQ_ ≥* 80.0). Additionally, an early-stop criterion is applied to avoid redundant computation under low-quality transformations. Specifically, the iteration is terminated if no new structures are generated or if *S_PQ_* fails to improve over three consecutive generations. Upon termination, the final population is re-ranked by IT-score and clustered. The top-10 models are selected for output.

### 4.7 Structure relaxation

Considering possible structural variations between the representative subunits and their corresponding structures in the subcomponents, the assembly process may yield models with steric clashes. Therefore, an extra step for structural relaxation is recommended to alleviate these clashes. This step effectively improves the quality of the models while not inducing significant structural alterations. In addition, the relaxation is also necessary for the IT-score scoring of assembled complex structures because IT-score is sensitive to severe atomic clashes. Structural relaxation is also advised during the sliding optimization step in the construction of cubic symmetric systems. In this study, all models generated by assembly process and O-symmetric docking approach are refined via 500 steps of molecular dynamics (MD) simulations using the ff14SB force field^69^ of AMBER (version 14)^70^.

### 4.8 Model scoring

The IT-score^50^, an iterative knowledge-based scoring function for protein-protein interactions, is used to score the final complexes in this study. Initially developed for evaluating pairwise interactions, IT-score is derived by iteratively optimizing interatomic pair potentials to maximize the discrimination between native and predicted structures based on the distributions of atom pairs, thereby circumventing the reference state problem in statistical potentials. For a dimer, the score is computed as a sum of the interatomic pair potentials within a cutoff of 10 A° between the receptor and ligand. In this work, we extend the IT-score to multimeric systems by decomposing the input complex into chain pairs and calculating IT-score for each pair. The final IT-score of a complex is obtained as the sum of the scores from all chain pairs. We utilize the IT-score to rank models generated by different methods, where a lower score indicates a higher structural quality.

### 4.9 Model clustering

To balance the result diversity and computational efficiency, different RMSD thresholds are applied at different clustering stages. A backbone RMSD of 5 A° is adopted during the local symmetry sampling stage in *C_n_* symmetric docking and *D_n_* symmetric docking stage. For the global sampling, assembly, and final model clustering stages, 5 A° *Cα* RMSD is used as the threshold.

A major challenge in clustering complexes with homologous subunits is identifying the optimal subunit correspondences. For a complex composed of *N* identical subunits, there are *N* ! equivalent permutations. Wrong subunit mappings may lead to high RMSD between similar models. Exhaustive enumeration of all possible mappings is computationally infeasible. To address this, a heuristic search strategy is adopted in this work^45^. Specifically, the centroids of each chain are calculated for both complexes. An initial superposition is performed by aligning the centroids in their original order. Then, for each pair of homologous subunits in the second complex, swap centroids and recalculate the centroid RMSD. If a swap leads to a lower RMSD, the new correspondence is retained. This process is iterated until no further RMSD reduction is observed. The final chain mapping is used for calculating the *Cα* RMSD between two complexes, thereby enhancing structural diversity in the final output.

### 4.10 Integration of cross-linking experimental data

We implement an extension to the assembly algorithm of HDM for allowing the integration of distance restraints derived from crosslinking experimental data. Specifically, after each assembly operation, we calculate the satisfaction ratio of crosslink-derived distance restraints associated with the subunits present in the newly formed subcomplex. A restraint is considered as being satisfied if the *Cα*-*Cα* distance between the corresponding residues is below the distance threshold specified by the restraint. For restraints involving repeated subunits, the restraint is regarded as being satisfied if at least one of the possible residue pairs meets the distance threshold. The satisfaction ratio of a subcomplex is defined as follows.

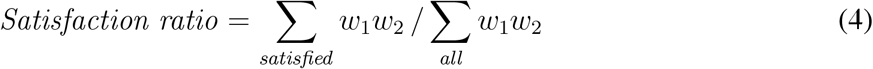

where *w*_1_ represents the experimental confidence of the crosslink restraint, such as the false discovery rate^57^, considering uncertainty in the experimental data. *w*_2_ represents the average pLDDT score of the residues involved in the satisfied restraint and is used to account for uncertainty in the predicted subunit structures. If the satisfaction ratio of a newly formed subcomplex falls below the predefined threshold, the corresponding assembly operation is discarded. During the initial sampling and genetic algorithm (GA) optimization stage, the satisfaction ratio threshold is set to 0.7 and 0.8, respectively. For the final full-complex models, both the confidence score and IT-score are multiplied by the satisfaction ratio to prioritize models fulfilling more distance restraints.

### 4.11 Reintegrating disordered terminal regions

To avoid interference with the docking process, the automated HDM pipeline includes a preprocessing step that removes disordered terminal regions. These trimmed regions can be reconstructed using approaches such as homology modelling, MD simulation, or structural alignment. As an illustration, we present the modelling and reintegration of terminal regions for the calcium/calmodulin-dependent protein kinase (PDB 1HKX) in Supplementary Fig. 9. Low-confidence disordered terminal residues are removed from the input monomer prior to docking (residues 1-6 and 143-147). HDM then performs symmetric docking on the filtered monomer, yielding a high-quality complex model (TM-score = 0.9702). The final full complex is obtained by aligning the full-length monomer onto each chain of the symmetric model using MMalign^68^.

### 4.12 Quality-weighted pairwise connectivity

The pairwise connectivity^45^ measures the maximum fraction of a full complex that can be assembled from the predicted subcomponents. Considering the impact of monomer quality on assembly performance, we introduce the quality-weighted pairwise connectivity (QPC), which explicitly account for the accuracy of extracted monomers. Specifically, each chain is treated as a node, with low-quality nodes filtered out based on monomer structural similarity (TM-score^71^ *<* 0.65 between the extracted and native monomers). Edges are then added between the remaining nodes if they are in contact within the target complex and a similar transformation (DockQ^72^ score *≥* 0.23) exists among the corresponding extracted transformations. After building the graph, we calculate its connected components. The QPC is defined as follows.

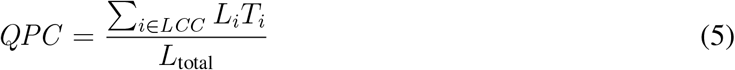

where *L_total_*denotes the total number of residues in the full complex, *LCC* represents the largest connected component, and *L_i_* and *T_i_* are the length and TM-score of the monomer corresponding to the node *i* in the *LCC*.

Overall, the QPC estimates the maximum potential to accurately assemble the full complex. Higher values indicate that a larger fraction of the complex could be accurately modelled, with a value of 1.0 indicating that the complete complex can potentially be assembled from the predicted subcomponents. Explicitly considering the accuracy of extracted monomers enables a better assessment of the quality of the structural information provided by the subcomponents for each target.

### 4.13 Accuracy assessment

We utilize two complementary metrics to evaluate the accuracy of predicted models, TM-score^71, 73^ for overall structural similarity and DockQ^72^ for the quality of interfaces between chains. The TM-score is calculated using the MMalign program^68^ after structural alignment between the predicted and native structures. Consistent with previous studies^42, 45^, a model is considered as acceptable-quality if its TM-score is above 0.7 and high-quality if TM-score is above 0.8. In addition, we also use the DockQ v2 program^54^ to compare predicted models with corresponding native structures and assess interface quality. Following the CAPRI criteria^74^, models are considered as acceptable-, medium-, and high-quality for 0.23 *≤ DockQ <* 0.49, 0.49 *≤ DockQ <* 0.80 and *DockQ ≥* 0.80, respectively. The success rate is defined as the fraction of benchmark targets yielding at least one acceptable-quality model within the top-1 predictions, unless otherwise specified.

### 4.14 Computational time

AF2 and AFM predictions are performed on NVIDIA A100 GPUs with 40 GB of memory. The runtime of AFM varies depending on the size of structures. For example, predicting a subcomponent comprising 2600 amino acids typically takes about 35000 s. Docking and assembly procedures are executed on 2.6 GHz Intel CPUs using multi-process parallelization, with 20 CPU cores employed by default. The average running times for the assembly method on two benchmarks are 5635 s and 5587 s, respectively. The docking stage generally consumes more time than the assembly process, particularly for heteromeric complexes. On Benchmark 2, the average running time for asymmetric docking is 10252 s, whereas symmetric docking takes an average of 4543 s. Overall, the running time for building a full complex structure can be reasonably finished within hours.

### 4.15 Comparison with other methods

Our HDM method is compared with two state-of-the-art structure prediction approaches for large protein complexes, CombFold^45^ and MoLPC^42^. For a fair comparison, the results of CombFold and MoLPC on Benchmark 1 and 2 are generated using the same subcomponent inputs as HDM, by running the open-source script from https://github.com/dina-lab3D/CombFold and https://github.com/patrickbryant1/MoLPC/. The results of AFM v2 on Benchmark 1 are obtained using our locally installed version. The AF3 predictions on Benchmark 2 and 3 are conducted using the AF3 server at https://alphafoldserver.com/. It is known that AlphaFold-predicted structures often contains hallucinating regions. Interestingly, the evaluation with only the regions of high quality results in slightly lower TM-scores, which is probably due to fewer amino acids to be aligned after removing low-confidence regions (Supplementary Fig. 10). Therefore, we used all amino acids of the predicted complex by AF3 during comparative evaluations. The results of other methods on Benchmark 3 are collected from the published paper^45^.

### 4.16 Visualizations

The images for all protein structures in this study were prepared using ChimeraX^75^.

## Data availability

All published data sets used in this paper were taken from the PDB (accession codes specified in the figure captions and in Supplementary Tables).

## Code availability

The HDM package is freely available at https://github.com/huang-laboratory/HDOCK-Multimer.

## Acknowledgements

This work was supported by the National Natural Science Foundation of China (grant Nos. 32430020, 32161133002, and 62072199) and the startup grant of Huazhong University of Science and Technology.

## Author contributions

S.H. conceived and supervised the project. X.Y. and Y.Y. designed and performed the experiments. X.Y., Y.Y., H.L., and S.H. analyzed the data. X.Y. and S.H. wrote the manuscript. All authors reviewed and approved the final version of the manuscript.

## Competing interests

The authors declare no competing interests.

## Notes

### Competing Interest Statement

The authors have declared no competing interest.

