## Supplementary material for "HDOCK-Multimer: integrating docking and combinatorial assembly for structure prediction of large protein complexes": Supplemtary Tables 1-4 and Figures 1-10

**Supplementary Table 1:** Chain and residue ranges of the divided groups used for HDM stepwise assembly on CASP15 target H1137.

| Chain | Group 1 | Group 2 | Group 3 |
|---|---|---|---|
| A | 1–173 | 153–316 | 296–409 |
| B | 1–156 | 136–300 | 280–343 |
| C | 1–155 | 135–298 | 278–524 |
| D | 1–165 | 145–316 | 296–547 |
| E | 1–174 | 154–320 | 300–390 |
| F | 1–157 | 137–300 | 280–518 |
| G / H | All | | |
| I | All | | |
| J | All | | |

**Supplementary Table 2:** Performances of HDM and other methods on 7 CASP15 targets of large protein complexes.

| | | | | | | Top-3 human predictors | | | Top-3 servers | | |
|---|---|---|---|---|---|---|---|---|---|---|---|
| Target | HDM | Comb Fold | MoLPC | AFM v2 | AF3 | Zheng | Venclovas | Wallner | Yang-Multimer | Manifold-E | MULTICOM_qa |
| H1111 | 0.91 | 0.80 | 0.52 | 0.08 | – | 0.98 | 0.98 | 0.09 | **0.98** | 0.98 | 0.94 |
| H1114 | 0.30 | – | 0.39 | 0.12 | – | 0.93 | **0.94** | 0.13 | 0.45 | 0.88 | 0.24 |
| H1137 | 0.78 | 0.87 | 0.60 | – | 0.59 | **0.94** | 0.82 | 0.64 | 0.67 | 0.88 | 0.76 |
| T1115 | 0.75 | 0.72 | 0.46 | – | **0.92** | 0.62 | 0.90 | – | 0.36 | 0.66 | 0.38 |
| T1165 | **0.81** | 0.79 | 0.75 | 0.75 | 0.79 | – | 0.77 | 0.76 | – | 0.80 | 0.78 |
| T1169 | **0.81** | **0.81** | 0.76 | 0.34 | 0.77 | – | 0.77 | 0.50 | – | 0.74 | 0.70 |
| T1192 | 0.87 | 0.86 | 0.88 | 0.50 | 0.81 | 0.88 | **0.89** | 0.83 | 0.88 | 0.88 | 0.89 |

**Supplementary Table 3:** Stoichiometry prediction results on 20 representative complexes from Benchmarks 1 and 2.

| PDB ID | Symmetry | Confidence score | Normalized IT-score |
|---|---|---|---|
| 6ZBB | Asymmetry | ✓ | |
| 6TDX | Asymmetry | | ✓ |
| 6RDB | Asymmetry | | |
| 7JG5 | Asymmetry | ✓ | |
| 5XPB | Helical | | |
| 1DPS | T | | |
| 2GRE | T | | ✓ |
| 2WYR | T | | ✓ |
| 6JCV | T | | ✓ |
| 6QLV | T | | ✓ |
| 1F1H | D6 | | ✓ |
| 4RAP | D6 | ✓ | ✓ |
| 5OVS | D7 | | ✓ |
| 2WCV | D5 | | ✓ |
| 2VYC | D5 | ✓ | ✓ |
| 1IJG | C12 | ✓ | ✓ |
| 2BL2 | C10 | ✓ | ✓ |
| 3TXQ | C11 | ✓ | |
| 2X2V | C13 | ✓ | ✓ |
| 4F4S | C10 | ✓ | ✓ |

**Supplementary Table 4:** HDM's performances on 4 antibody–antigen complexes from Benchmarks 1 and 2.

| PDB ID | HDM TM-score | | HDM DockQ | |
|---|---|---|---|---|
| | Top 1 | Top 10 | Top1 | Top10 |
| 2VYR | 0.2597 | 0.2597 | 0.020 | 0.026 |
| 6SSI | 0.8739 | 0.9322 | 0.559 | 0.669 |
| 6X04 | 0.3052 | 0.3707 | 0.017 | 0.042 |
| 4RDQ | 0.8639 | 0.8639 | 0.263 | 0.330 |

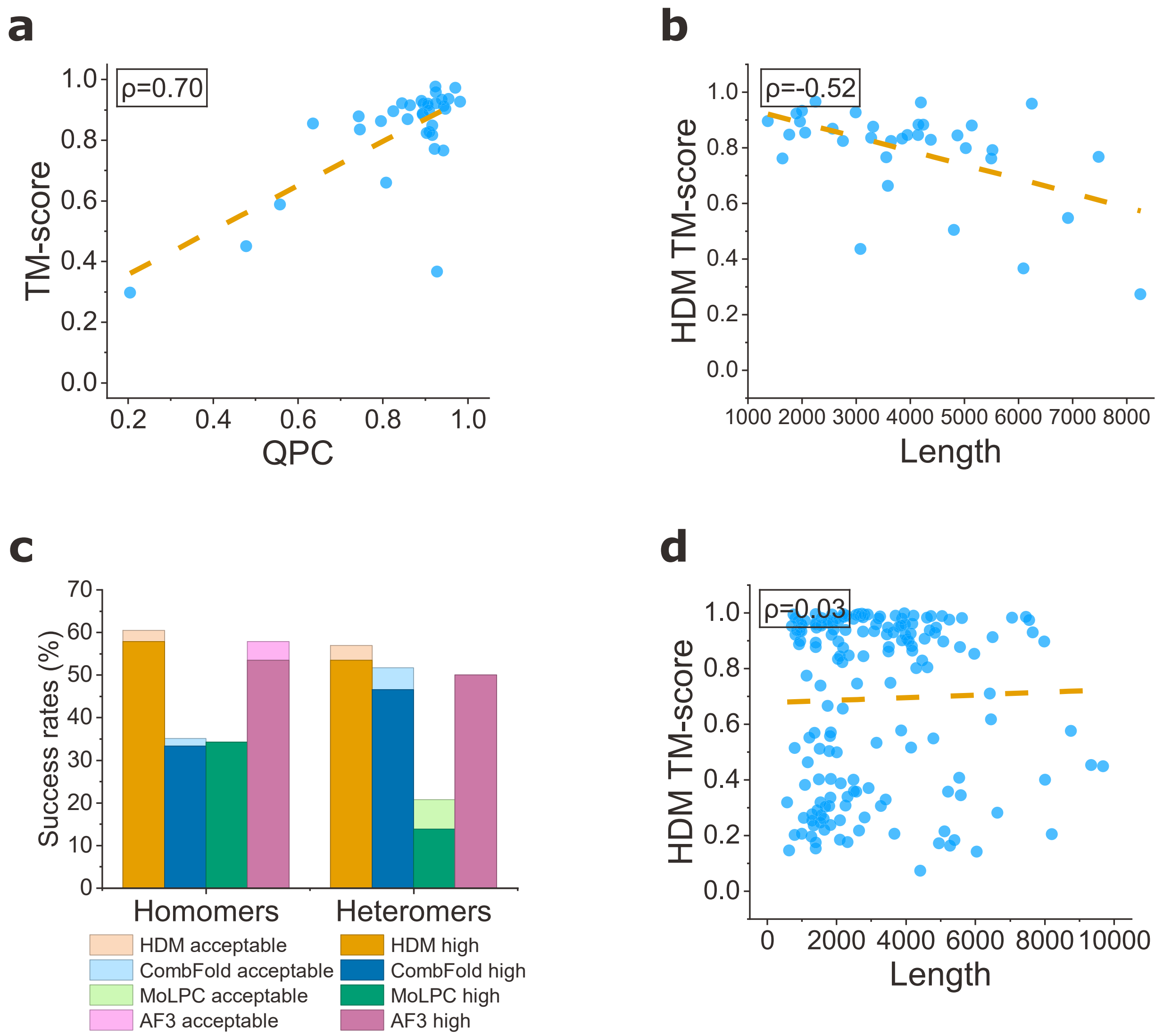


**Supplementary Fig. 1: Analysis of impacting factors on HDM performance. a**, Top-10 TM-score as a function of Quality-weighted pairwise connectivity (QPC) on Benchmark 1. **b**, Top-1 TM-score as a function of complex size (number of amino acids) on Benchmark 1. **c**, Top-1 success rates for homomeric and heteromeric complexes on Benchmark 2. **d**, Top-1 TM-score as a function of complex size (number of amino acids) on Benchmark 2.

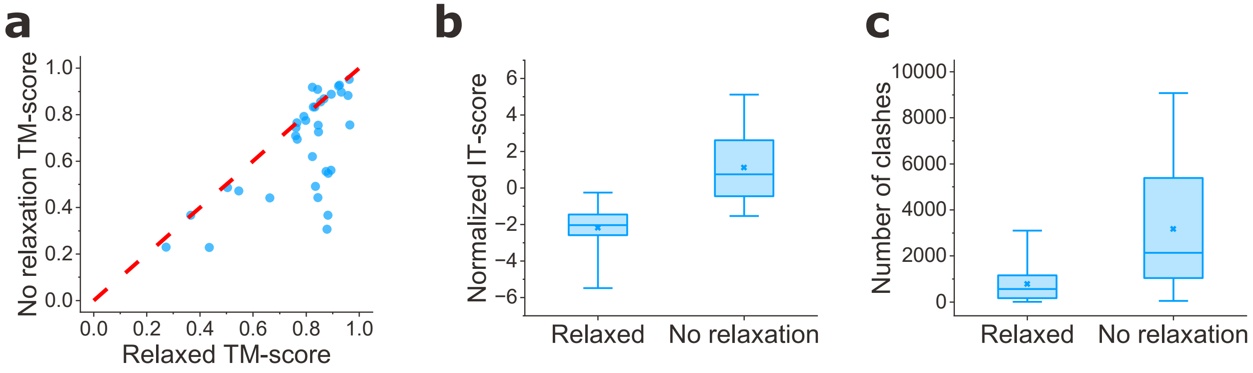


**Supplementary Fig. 2: Performances of HDM before and after MD relaxation on Benchmark 1.** **a**, Head-to-Head TM-score comparison for the models predicted by HDM before and after relaxation. **b**, Distributions of the normalized IT-scores for the models predicted by HDM before and after relaxation. **c**, Distributions of the number of atomic clashes for the models predicted by HDM before and after relaxation.

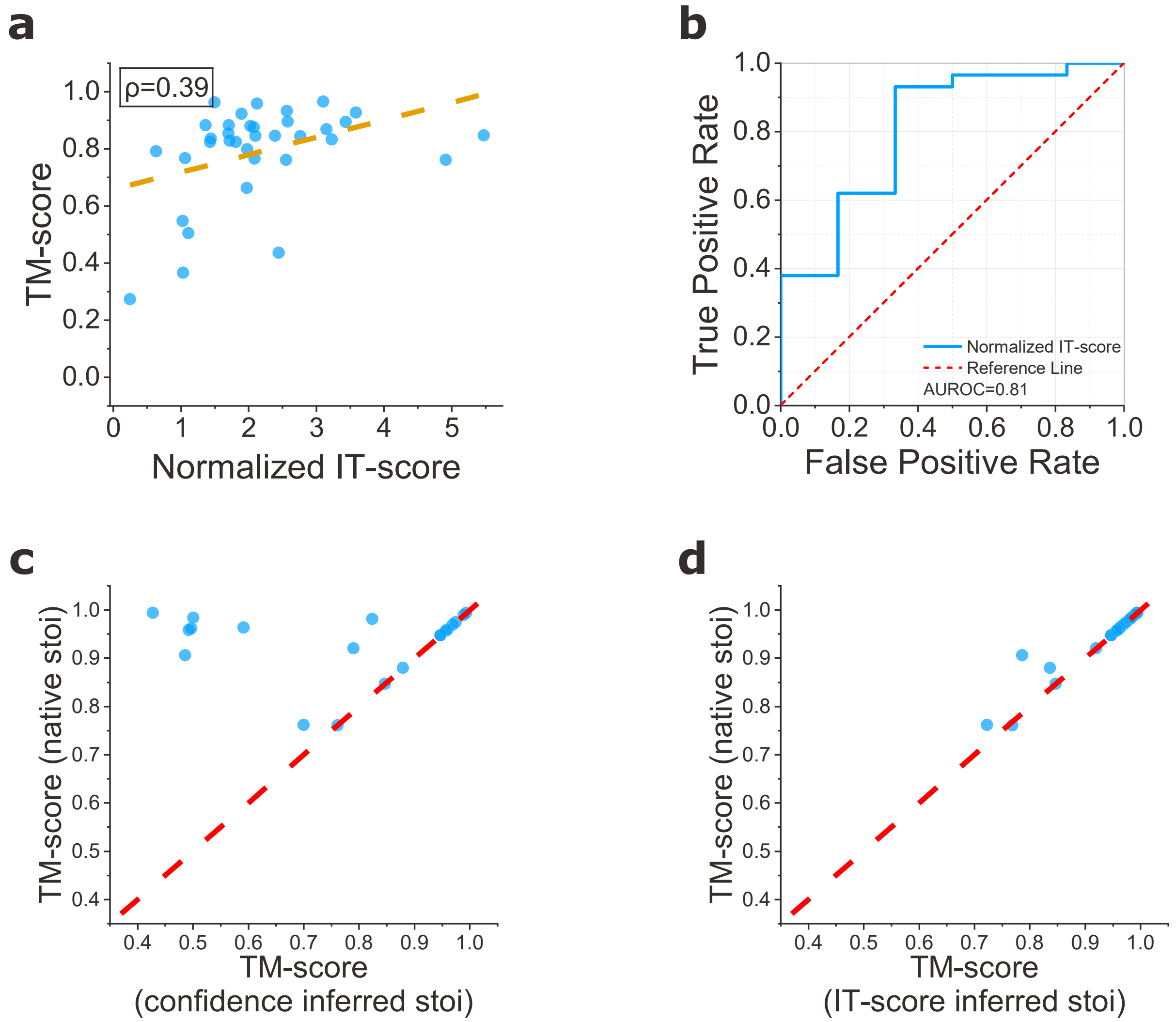


**Supplementary Fig. 3: Stoichiometry prediction.** **a**, Top-1 TM-score versus normalized IT-score on Benchmark 1. The normalized IT-score is negated to yield a positive correlation. **b**, Receiver operating characteristic (ROC) curves assessing the ability of the normalized IT-score to distinguish successful predictions (TM-score $> 0.7$) on Benchmark 1. **c**, TM-score comparison of HDM with native stoichiometries and stoichiometries inferred from confidence scores. **d**, TM-score comparison of HDM with native stoichiometries and stoichiometries inferred from normalized IT-scores.

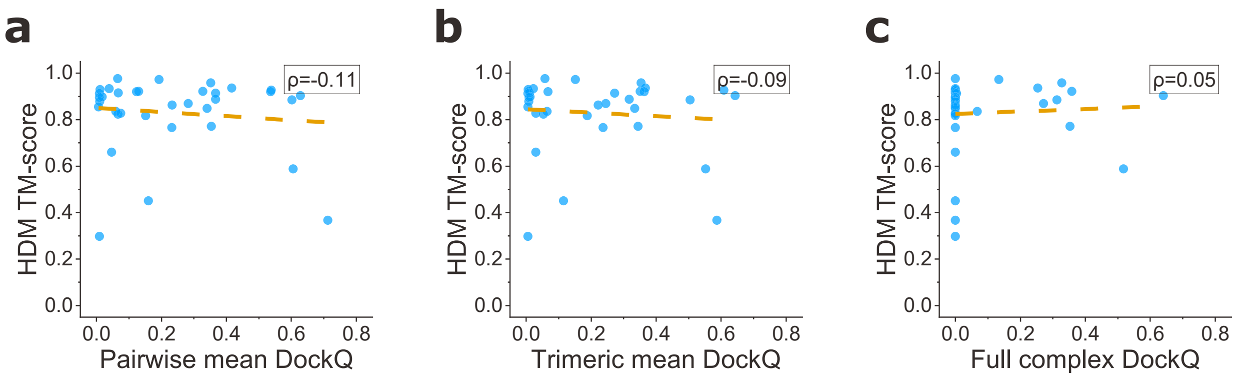


**Supplementary Fig. 4: HDM performances as a function of interfaces novelty on Benchmark 1.** **a**, Top-10 TM-score versus mean DockQ for all pairwise interfaces in the native structure. **b**, Top-10 TM-score versus mean DockQ for all trimeric interfaces in the native structure. **c**, Top-10 TM-score versus DockQ for the full complex.

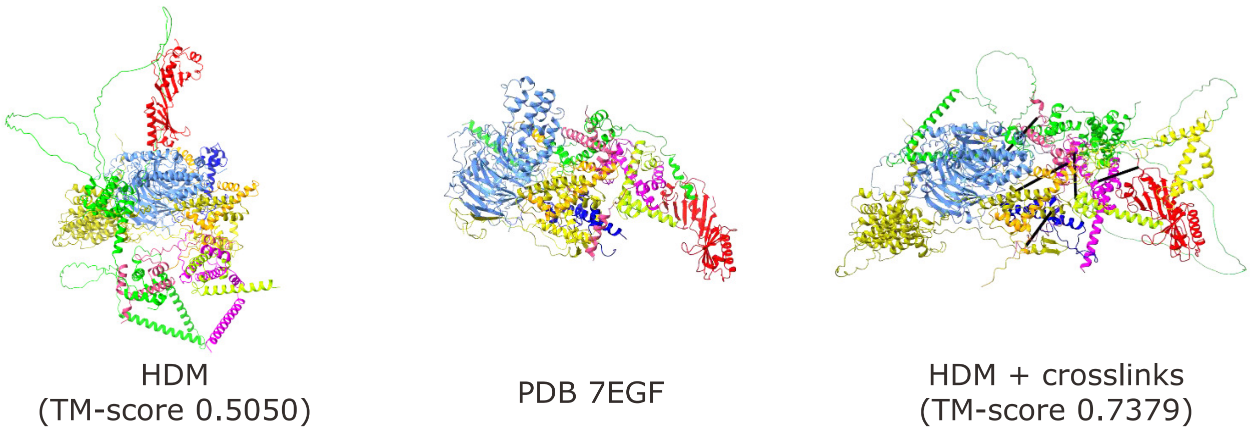


**Supplementary Fig. 5: Improvement of HDM prediction for the TFIID lobe A subcomplex (PDB 7EGF) using crosslink restraints.** From left to right are inaccurate HDM model without crosslink restraints (left), cryo-EM structure (middle), and acceptable-quality model by HDM with crosslink restraints (right). Crosslinks are shown as black lines.

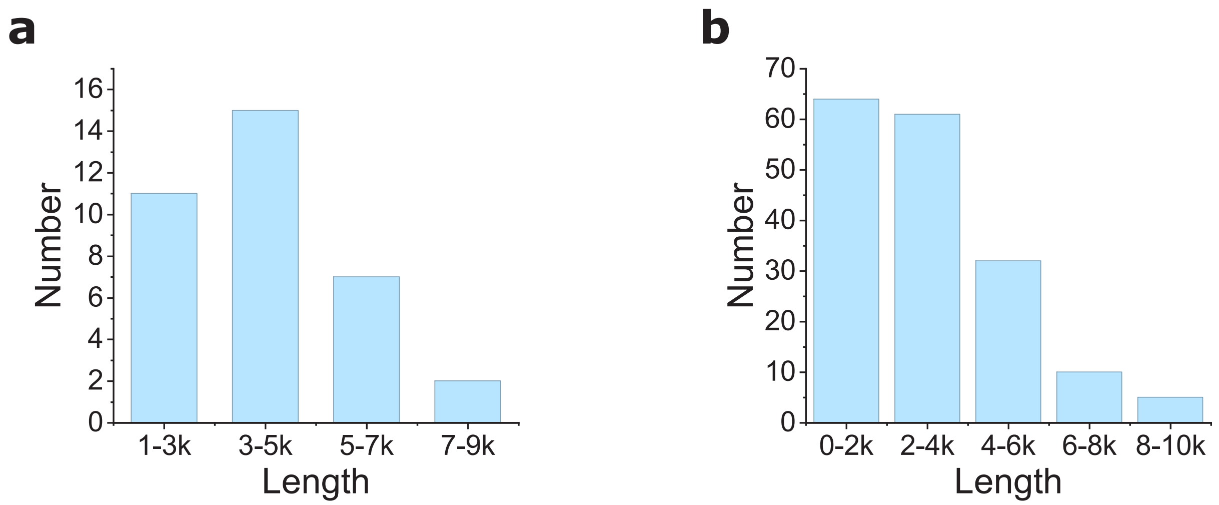


**Supplementary Fig. 6: Length distribution of the targets in Benchmark 1 and 2.** Here, the length stands for the total number of amino acids in a complex. **a**, Length distribution of the complexes in Benchmark 1. **b**, Length distribution of the complexes in Benchmark 2.

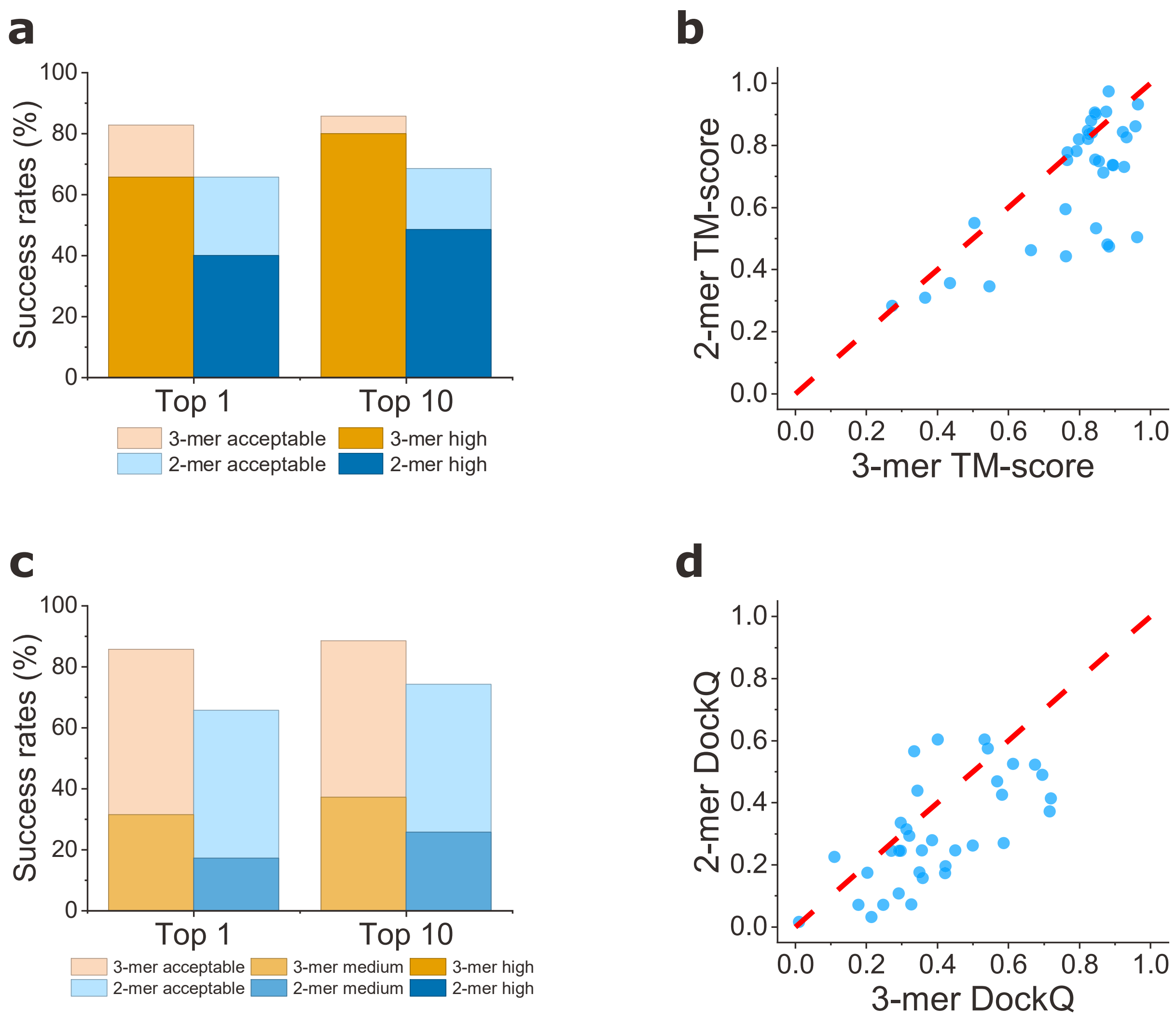


**Supplementary Fig. 7: Effect of subcomponent size on HDM performance on Benchmark 1. a**, Comparison of the average success rates of HDM using dimers or trimers, evaluated by TM-score. **b**, Head-to-head TM-score comparison for the models predicted by HDM using dimers or trimers. **c**, Comparison of the average success rates of HDM using dimers or trimers, evaluated by DockQ. **d**, Head-to-head DockQ comparison for the models predicted by HDM using dimers or trimers.

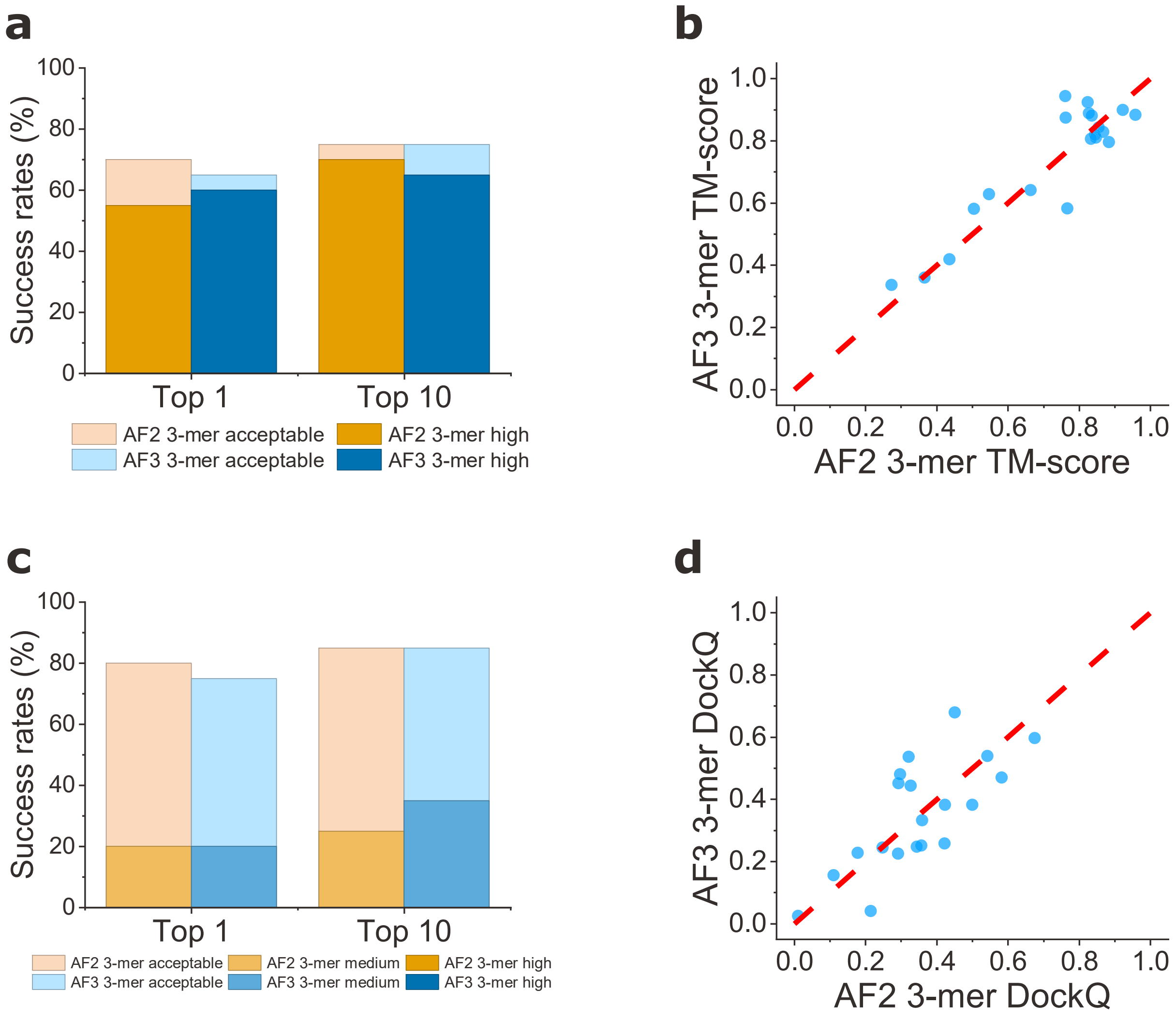


**Supplementary Fig. 8: Effect of subcomponent source on HDM performance on a Benchmark 1 subset (n = 20). a**, Comparison of the average success rates of HDM using AF2- or AF3-predicted trimers, evaluated by TM-score. **b**, Head-to-head TM-score comparison for the models predicted by HDM assembly using AF2- or AF3-predicted trimers. **c**, Comparison of the average success rates of HDM using AF2- or AF3-predicted trimers, evaluated by DockQ. **d**, Head-to-head DockQ comparison for the models predicted by HDM assembly using AF2- or AF3-predicted trimers.

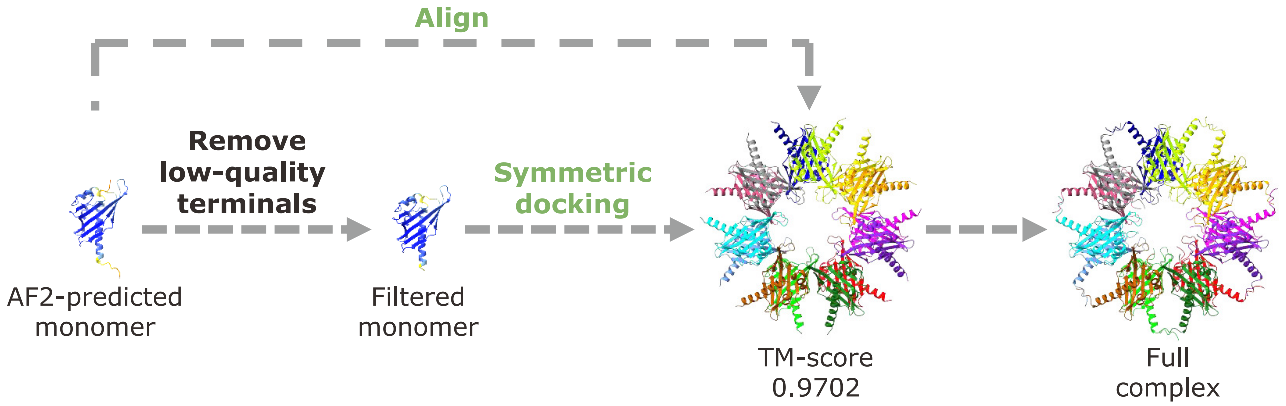


**Supplementary Fig. 9: An illustration of reintegrating disordered terminal regions for target 1HKX.** Specifically, low-confidence disordered terminal residues are removed from the input monomer prior to docking (residues 1-6 and 143-147). HDM then performs symmetric docking on the filtered monomer, yielding a high-quality complex model (TM-score = 0.9702). The final full complex is obtained by aligning the full-length monomer onto each chain of the symmetric model using MMalign.

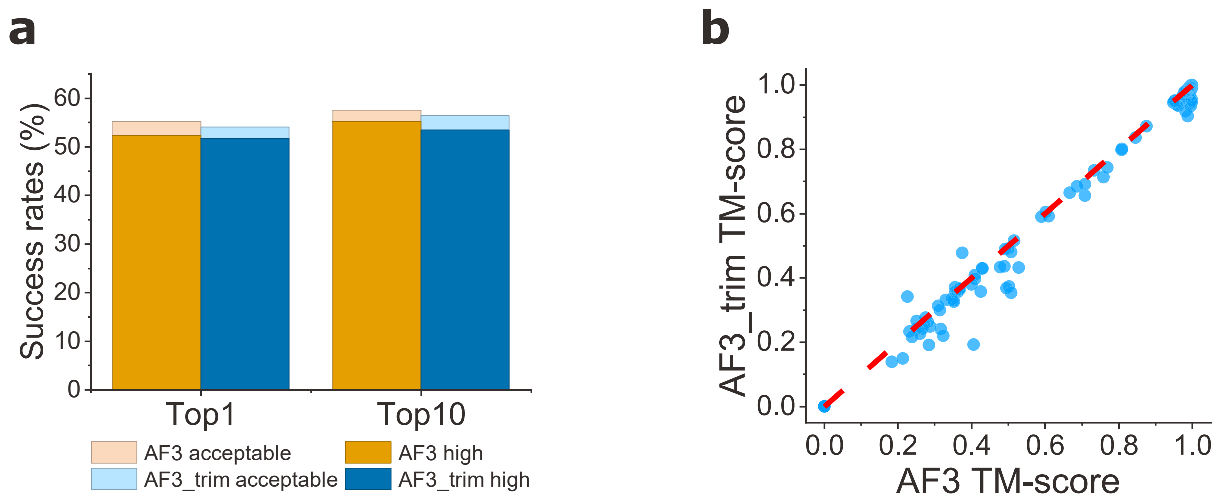


**Supplementary Fig. 10: Effect of trimming low-confidence regions on AF3 model accuracy.** **a**, Comparison of the average success rates of AF3 before and after trimming low-confidence regions (pLDDT<70) on Benchmark 2, evaluated by TM-score. **b**, Head-to-head TM-score comparison for the AF3-predicted models before and after trimming low-confidence regions on Benchmark 2. Trimming was performed using the same criteria as in monomer preprocessing.
